# Damaraland mole-rats are not obligate cooperative breeders

**DOI:** 10.1101/2021.12.08.471794

**Authors:** Jack Thorley, Hanna M. Bensch, Kyle Finn, Tim Clutton-Brock, Markus Zöttl

## Abstract

Damaraland mole-rats (*Fukomys damarensis)* are usually viewed as a eusocial or obligate cooperative breeder in which successful reproduction is dependent on help from closely related group members. However, because longitudinal studies of mole-rats in their natural environment are uncommon, the extent to which successful reproduction by breeders relies on assistance from other group members is unclear, and for non-breeders, the immediate and delayed fitness consequences of staying and helping are poorly understood. Using data from a 7-year field study of marked individuals, we investigate whether increasing group size confers reproductive and survival benefits to breeders and non-breeders, and explore the different routes through which individuals acquire breeding positions. We show that solitary natal dispersal was the most common route to breeding for both sexes and that the inheritance of dominant breeding positions was uncommon in both sexes. After dispersing, females typically settled alone in new burrow systems where they enjoyed high survival rates and remained in good body condition - often for several years - before being joined by males. In contrast to most obligately cooperative species, pairs of potential breeders reproduced successfully without helpers and experimentally formed pairs had the same reproductive success as larger established groups. Though larger breeding groups recruited slightly more pups on average, our data suggest that neither survival nor reproduction depend on the presence of non-breeding helpers, indicating that Damaraland mole-rats are not obligate cooperative breeders. We suggest that extended philopatry and group living in Damaraland mole-rats have evolved because of the high costs and constraints of dispersal rather than because of strong indirect benefits accrued through cooperative behaviour and that similarities between their breeding systems and those of obligatorily eusocial insects have been over-emphasized.

**Significance Statement:** The social mole-rats are often considered eusocial mammals in which successful reproduction depends on assistance from non-breeding helpers. In this study we show that in wild Damaraland mole-rats, the presence of non-breeders is associated with both costs and benefits and that nascent breeding pairs show high reproductive success despite the lack of non-breeding helpers. These findings indicate that Damaraland mole-rats are not obligate cooperative breeders and suggest that similarities between their breeding systems and those of obligatorily eusocial insects have been over-emphasized.

## INTRODUCTION

In some animal societies juveniles choose to remain in their natal group for extended periods of time and do not breed instead of dispersing and attempting to breed independently (1–6). In some of these species, retained offspring but provide no direct assistance in rearing young produced by dominant breeders (7-10), while in others that are usually referred to as cooperative breeders, they provision young born in the group (3, 4, 11–14). In ‘facultative’ cooperative breeders, breeding groups may or may not include non-breeding helpers and breeders are able to rear young without assistance from non-breeders while in more specialised ‘obligate’ cooperative breeders, helpers have substantial effects on the reproductive success of breeding individuals and breeders are usually unable to rear young successfully without assistance from non-breeders (4, 9, 15–17).

The social African mole-rats (*Bathyergidae*), which include naked mole-rats (*Heterocephalus glaber*) and Damaraland mole-rats (*Fukomys damarensis*), are often regarded as among the most specialised cooperative breeders (18-20). Groups of social mole-rats usually consist of a single breeding pair and several litters of offspring that have delayed dispersal and refrain from reproduction while living in their natal group (21-28). The principal cooperative activity of non-breeding individuals is to maintain a network of tunnels that provide access to underground plant tubers and to bring food to food stores that are accessible to all group members (29-31), though non-breeders of both sexes also groom and retrieve pups born to the dominant female if they stray from the breeding nest (30, 32–34).

The extensive involvement of many non-breeders in cooperative foraging has led to the suggestion that social mole-rats rely on a workforce for reproduction and survival, and the idea that there are strong benefits of group-living to breeders is implicit in most discussions surrounding the evolution of sociality in the African mole-rats (35-39). Early studies of mole-rats emphasized similarities between their breeding systems and those of obligatorily eusocial insects (19,40,41) and both naked and Damaraland mole-rats are often referred to as eusocial (28, 29, 36, 38, 42–45). However, despite the broad interest in the evolution of mole-rat sociality, longitudinal studies of wild mole-rat populations remain uncommon (24,28,46,47). One study of wild Damaraland mole-rats has investigated the demographic effects of group size in social mole-rats (39) and has shown that larger groups recruited more offspring. In the same study, juvenile survival was independent of group size and juvenile growth rate was reduced in large groups, possibly by competition between non-breeders. Breeding females have also been shown to have lower workloads than non-breeders, both in a separate population of Damaraland mole-rats (48) and in captive animals (49). It is not yet known whether breeding pairs without helpers are able to raise offspring to independence, and it is still unclear whether social mole-rats are facultative or obligate cooperative breeders and whether living in groups increases the survival of group members.

We also know little about the dynamics of mole-rat groups or about the paths by which individual achieve breeding status in natural mole-rat populations. In some of the most well-studied obligate cooperative mammals, pairs of individuals without helpers usually fail to breed successfully (17,50,51). New groups typically form and successfully breed when coalitions of individuals disperse and settle together (52, 53), when large groups fission into smaller, distinct breeding units (54-57), or when a breeding territory including helpers is inherited by existing group members (58-61). Similar patterns of group formation as in obligate cooperative mammals are seen in many eusocial insect societies (62-65), and various studies have suggested that the fissioning of large groups may also be an important driver of new group formation in the social mole-rats (19,21,31,66). However, research on numerous mole-rat populations also indicates that the dispersal of solitary individuals is widespread (24, 41, 66–68), and evidence of individuals living singly in burrow systems has repeatedly been remarked upon (24,68,69). The survival rate of single individuals and the frequency with which these individuals form new breeding groups has not been quantified, but if it represents an important route to breeding, then it would suggest that mole-rats do not need a workforce to reproduce successfully.

In this study we investigate how new breeding groups emerge and determine whether reproductive success depends on the presence of non-breeders in a wild population of Damaraland mole-rats. We first describe how new groups form and then we assess whether individuals living in groups (breeders and non-breeders) experience higher survivorship than individuals that have dispersed and settled solitarily. By experimentally creating breeding pairs in the field, we examine whether the reproductive output of breeders lacking non-breeding group members is lower than that of established groups which have access to a potential workforce. Lastly, we use demographic information collected across 7 years of field study to analyse the associations of group size with (i) reproductive success, (ii) adult survival, and (iii) growth of recruited offspring, testing the prediction that large group size is associated with increases in reproductive success and survival, and enhances growth among young individuals.

## RESULTS

### a) Population structure and distribution of group sizes

The mean group size at our study site in the Kalahari across all captures between 2013 and 2020 was 5.91 ± 5.61 individuals (mean ± SD; max = 26; n = 328 group captures; Fig. 1). Of the 84 unique groups that were captured, 49 were of a single individual on the first capture, 94% of which were females (n = 46). Excluding this large number of single individuals, the mean group size was 8.67 ± 5.30 individuals (mean ± SD; n = 210 group captures), which can be taken to represent the average size of a breeding unit in the population. Across the duration of the study only one reproductively active ‘breeding’ female was present at a group at any given time.

**Fig. 1.**
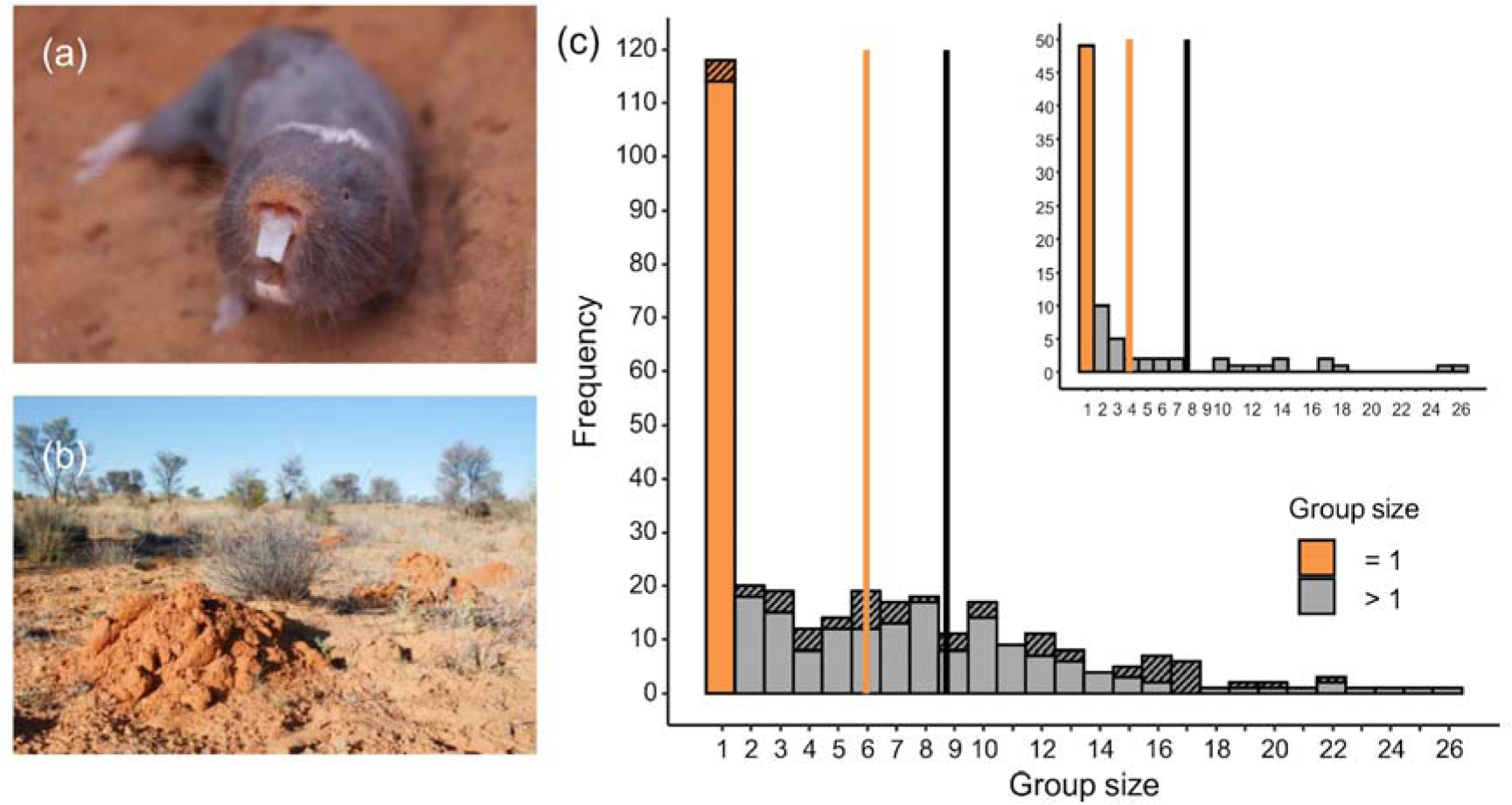
Group sizes for Damaraland mole-rats in the southern Kalahari. (a) An adult Damaraland mole-rat. (b) A line of sand mounds that is created by a group of mole-rats when they dig foraging tunnels underground. (c) The distribution of group sizes in the study population at the Kuruman River Reserve. The main histogram displays group size frequencies across all captures, while the inset histogram displays the group size at first capture for each unique group (n = 82; including single individuals). In each case, a large proportion of the population were captured as single individuals (orange), who were almost exclusively females. The vertical lines display the mean group size either including (orange) or excluding (black) these single individuals, and for all captures, we distinguish between complete and incomplete group captures by hatching the latter (mean for all complete group captures = 5.45, 1SD = 5.51; mean for complete captures of groups > 1 = 8.54, 1SD = 5.32). Removing the 15.2% of incomplete captures had limited influence on the mean group size (not displayed).

### b) Dispersal and routes to breeding

Males and females dispersed from their natal group at a similar age (Fig. S8; MSM; hazard ratio of males relative to females = 1.125, 95% CI = [0.864,1.476]; see SI for philopatry analyses), approximately 1.80 years after their first capture for males (95% CI [1.50,2.15]), and 2.02 years after their first capture for females (95% CI [1.66, 2.47]). The timing of departure from the natal group coincided with periods when ecological constraints were relaxed through increased rainfall (MSM: hazard ratio = 1.355, 95% CI [1.205,1.523]; Table S1).

The large number of single females reflects a broader divergence in the life history trajectories of the two sexes after departure from their natal group. Post-dispersal, females usually settled as single individuals, whereas males left their natal group and sought to locate females and seldom settled on their own. We identified 23 females in our population whose route to breeding could be determined accurately. Of these 23, 60.9% (n = 14) were females that had dispersed from their group, settled singly, and were later joined by one or more emigrant males to start a nascent breeding group. Two further females became breeders through territory budding, having dispersed and subsisting alone in a burrow system directly adjacent to but separate from their natal burrow, with males subsequently immigrating in. In contrast, 30.3% of breeding females (n = 7) inherited a breeding position in their natal burrow system after the disappearance of the previous dominant female. Females never immigrated into established groups and cases of female inheritance were always associated with the loss of the incumbent breeding female, either following group collapses that left a single female that was later joined by a male forming a new pair (n = 3), or replacement of the breeding female by a natal female who continued to reproduce in the group (n = 4).

For males, joining single females in newly established or inherited burrow systems (n = 16), or immigrating into established groups (n = 13), were both common routes through which males accessed unfamiliar females. Without paternity information it remains unknown whether these males were the father of all offspring born post-immigration; though cases of multiple male immigration (from either the same or from different groups) were infrequent (n = 3 cases). For the same reason, it is unclear whether males ever inherited breeding positions in their natal group; however, the fact that groups often collapsed after the death of the breeding female, coupled with the lack of female immigration, makes male inheritance improbable.

### c) Status-related survivorship among females

Once settled, single females exhibited high annual survival rates and disappeared from the population at rates intermediate between those of breeders and those of non-breeders (Fig. 2, Table S4); their estimated annual survival was higher (79.3%) than that of in-group non-breeding females (64.2%) and was lower than that of breeding females (86.7%). Thus, in-group non-breeding females were 1.91 times more likely to disappear from the population than single females (MSM: ratio of transition probabilities; 95% CI [1.07,3.43]), and 3.28 times more likely to disappear than breeding females (MSM: 95% CI [1.84,5.82]). The rate of disappearance of single females was higher than that of breeding females, with the former being 1.71 times more likely to disappear than the latter, though this effect was not significant (MSM: 95% CI [0.78,3.77], Fig. 2c). Group size did not affect the likelihood that non-breeding females (MSM: hazard ratio = 1.003, 95% CI [0.973,1.033]) or breeding females (MSM: hazard ratio = 1.058, 95% CI [0.970,1.156]) disappeared.

**Fig. 2.**
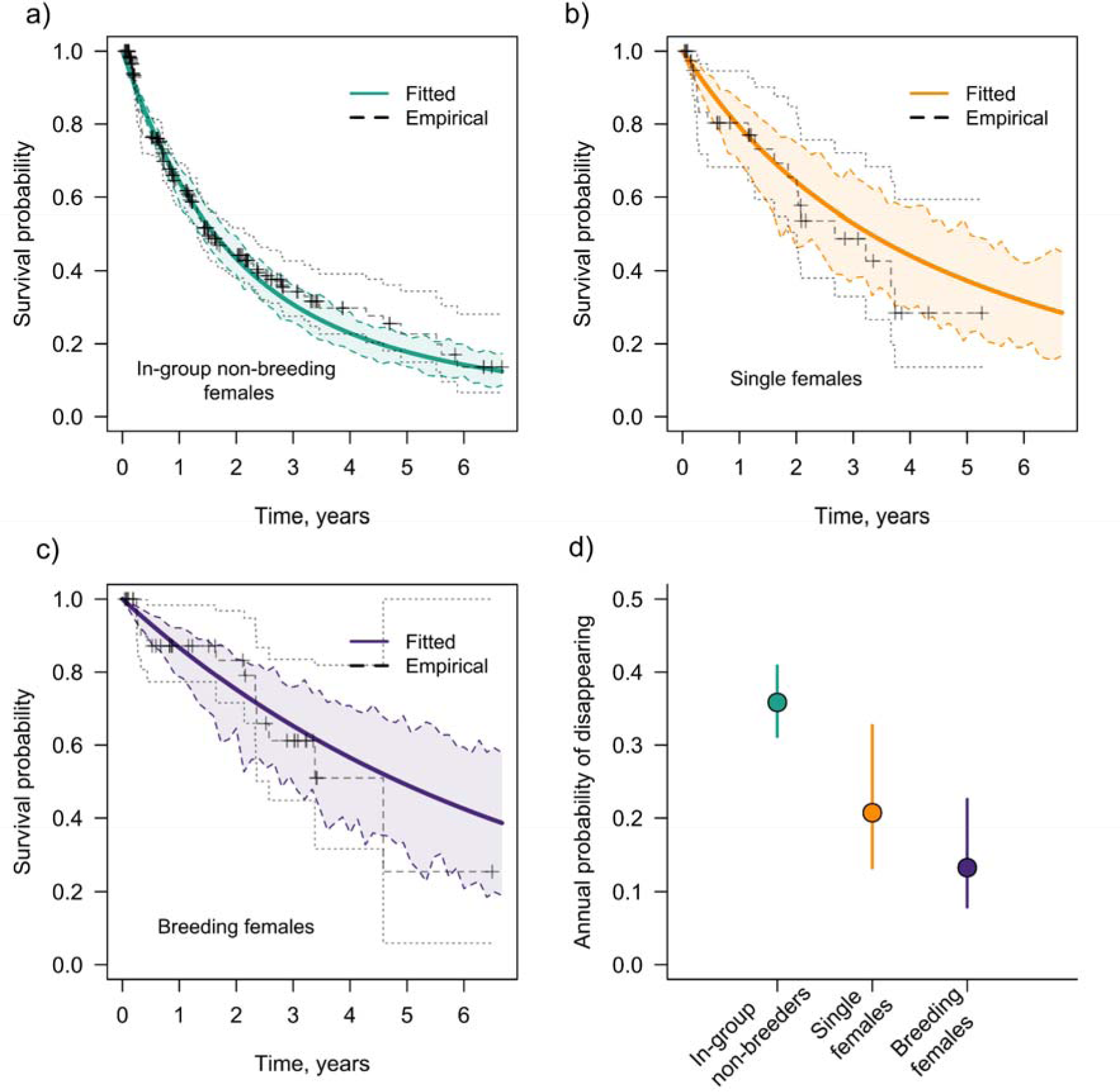
Status-related survivorship of female Damaraland mole-rats: in-group non-breeders (a), single females (b), and breeding females (c). Solid lines display the expected probability of survival from each state as estimated from the multi-state Markov model (see methods). Dotted lines display the ‘empirical’ Kaplan-Meier estimate of survival probability, with crosses denoting cases of right-censoring. Here, survival strictly refers to disappearance from the study population, and the predicted annual probability of disappearance (mean ± 95% CI) is given in (d). ‘Survival’ therefore combines cases of in-state mortality with cases of dispersal, where individuals have left their current group and state and are not re-captured thereafter. Dispersal is most likely to account for a large proportion of disappearances from the in-group non-breeder state (Table S1), and the low incidence of recaptures of individuals thereafter implies that dispersal carries a high cost of mortality. The multi-state model combined information on 326 *in-group non-breeding females,* 45 *single females*, and 46 *breeding females* (though females could appear in multiple states). Note that one female had been recaptured as a single individual for 5.26 years by the end-point of the study, and was right-censored at that point.

Single females maintained good body condition over extended periods, frequently over several years. Their body condition was equivalent to that of size-matched non-breeding females (Fig. 3), suggesting that solitary living does not affect the ability of individuals to acquire food and maintain mass (condition analyses outlined in SI).

**Fig. 3.**
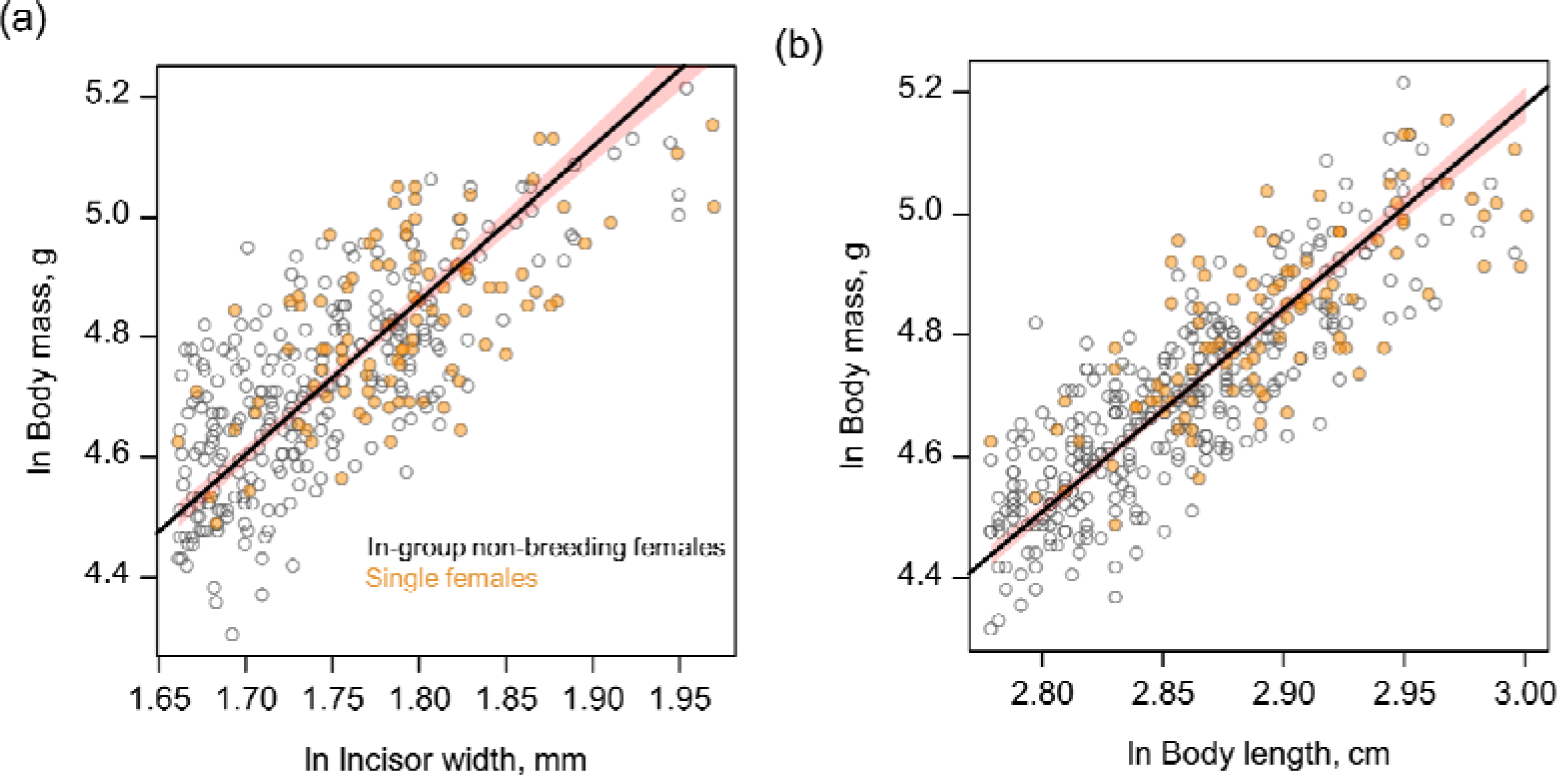
The body condition of dispersed single females compared to size-matched in-group non-breeding females. The principal plots display the scaled major axis regression (SMA) between body mass and incisor width (a), and between body mass and body length (b). Points display the raw data and lines display the predicted regression through all the data, with 95% bootstrapped confidence intervals given by the shaded area. Allowing the two classes of females to have different intercepts and slopes did not improve model fit: the two classes of females did not differ in mean body condition, or in the change in body condition with increasing skeletal size. Incisor width data (n = 353) includes 255 measures from in-group non-breeding females (n = 151 unique individuals) and 98 from single females (n = 44 unique individuals). Body length data (n = 419) includes 320 measures from in-group non-breeding females (n = 187 unique individuals) and 99 from single females (n = 44 unique individuals)

### d) Within-group recruitment

Experimentally created pairs recruited a similar number of offspring into their group as established groups that were captured and recaptured over the same time period (Fig. 4a; Welch’s t-test, t = 0.253, df = 14.02, p = 0.80; Fig. S8 details capture history after pairing). The recruitment rate in experimental and control groups over the experimental period was comparable to the total recruitment rate over the whole study period (Fig. 4a), and environmental conditions across the experimental period were typical of those seen in the Kalahari over a longer time span (Fig. S10).

**Fig. 4.**
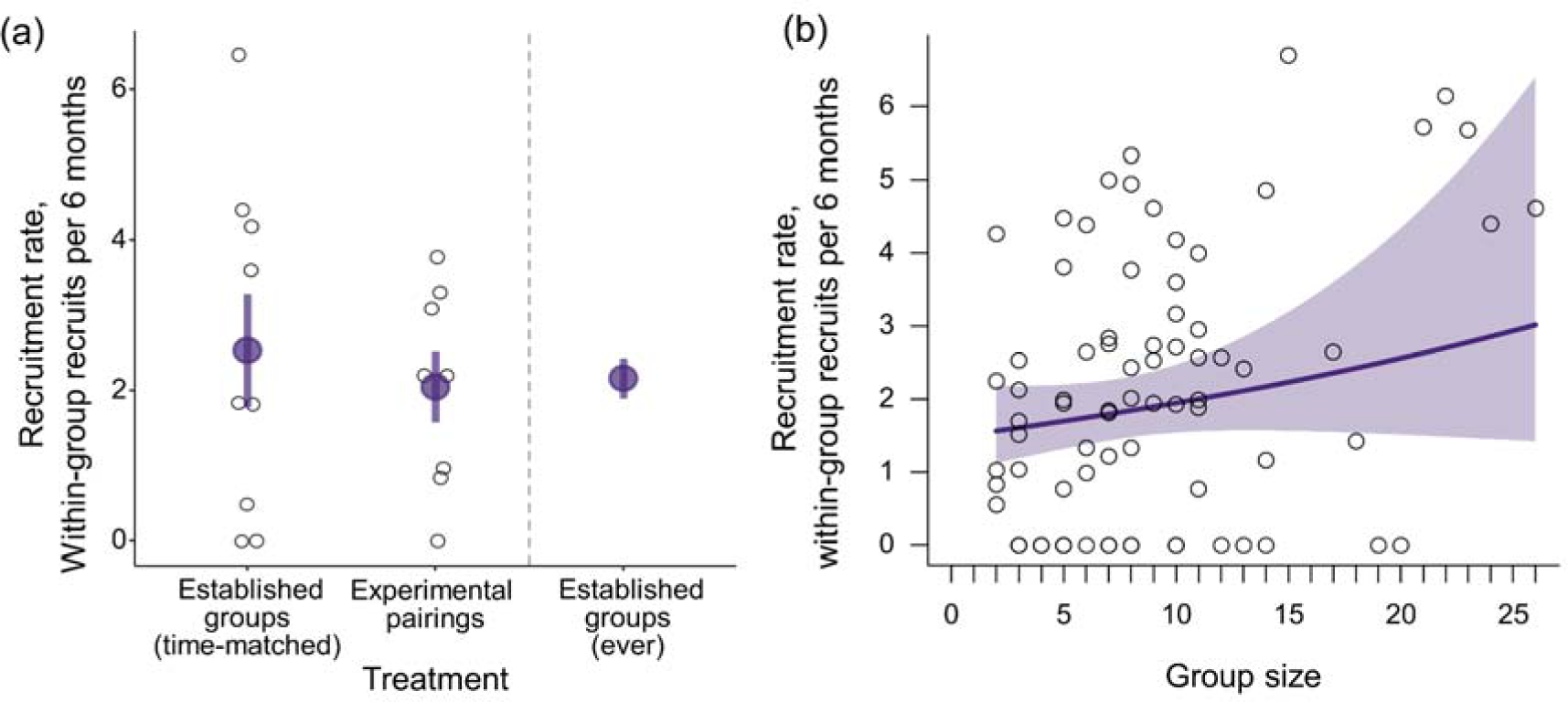
The effect of group size on within-group recruitment. a) The recruitment rate of established groups (n = 9) did not differ from that of newly created pairs (n = 8) when measured over the same time period. The mean rate of recruitment in this experimental period (middle panel in 5a) was similar to the overall mean recruitment rate in the population (right panel in 5a; average of all values in 5b). The larger purple points denote the treatment mean ± 1 SEM. b) In contrast, longitudinal analyses of group size (n = 78) across the duration of study detected a modest effect of increasing group size on the rate of within-group recruitment (0.216 ± 0.106, z = 2.04, p = 0.042, GLMM, link-scale); solid line shows the predicted mean and shading shows the 95% confidence intervals. In both panels the black unfilled points give raw data, corrected for the trapping interval duration, and in all cases, recruitment has been standardised to a 6-monthly rate according to the time difference between the first capture and the second capture.

Across the duration of the study, increasing group size was associated with a modest increase in within-group recruitment rate (GLMM: 0.216 ± 0.106, z = 2.04, p = 0.042, Table S4, Fig. 4b). Changing the group size term to reflect the mean group size across the capture and recapture event did not qualitatively affect the result. Increased rainfall in the year prior to the first trapping showed a positive but non-significant effect on recruitment (geometric mean monthly rainfall, GLMM: 0.159 ± 0.086, z = 1.85, p = 0.065), and the body mass of the breeding female did not significantly affect recruitment (GLMM: -0.090 ± 0.089, z = -1.02, p = 0.31). The inclusion of an interaction between rainfall and group size suggested that the effect of group size on recruitment rate did not depend upon recently experienced weather conditions (GLMM: 0.108 ± 0.080, z = 1.36, p = 0.17).

### e) Early-life growth and adult body mass

In both sexes, increases in group size were associated with faster growth rates in early life (Fig. 5, Table S5). However, in larger groups, growth rates declined more strongly during ontogeny and their members ultimately went on to attain a lower asymptotic body mass (NLMM: male *A_GS_* = -8.10 ± 2.02, p < 0.001; female *A_GS_* = -9.42 ± 1.47, p < 0.001; Fig. S11, see SI for further treatment). Increases in group size were also associated with reductions in asymptotic skeletal size in both sexes, though the effect was only supported in females (NLMM: male *A_GS_* = -0.047 ± 0.036, p = 0.19; female *A_GS_* = -0.113 ± 0.027, p < 0.001, Table S7). In contrast to the body mass models, there was no support for a relationship between group size and the growth rate constant in either sex (Table S7).

**Fig. 5.**
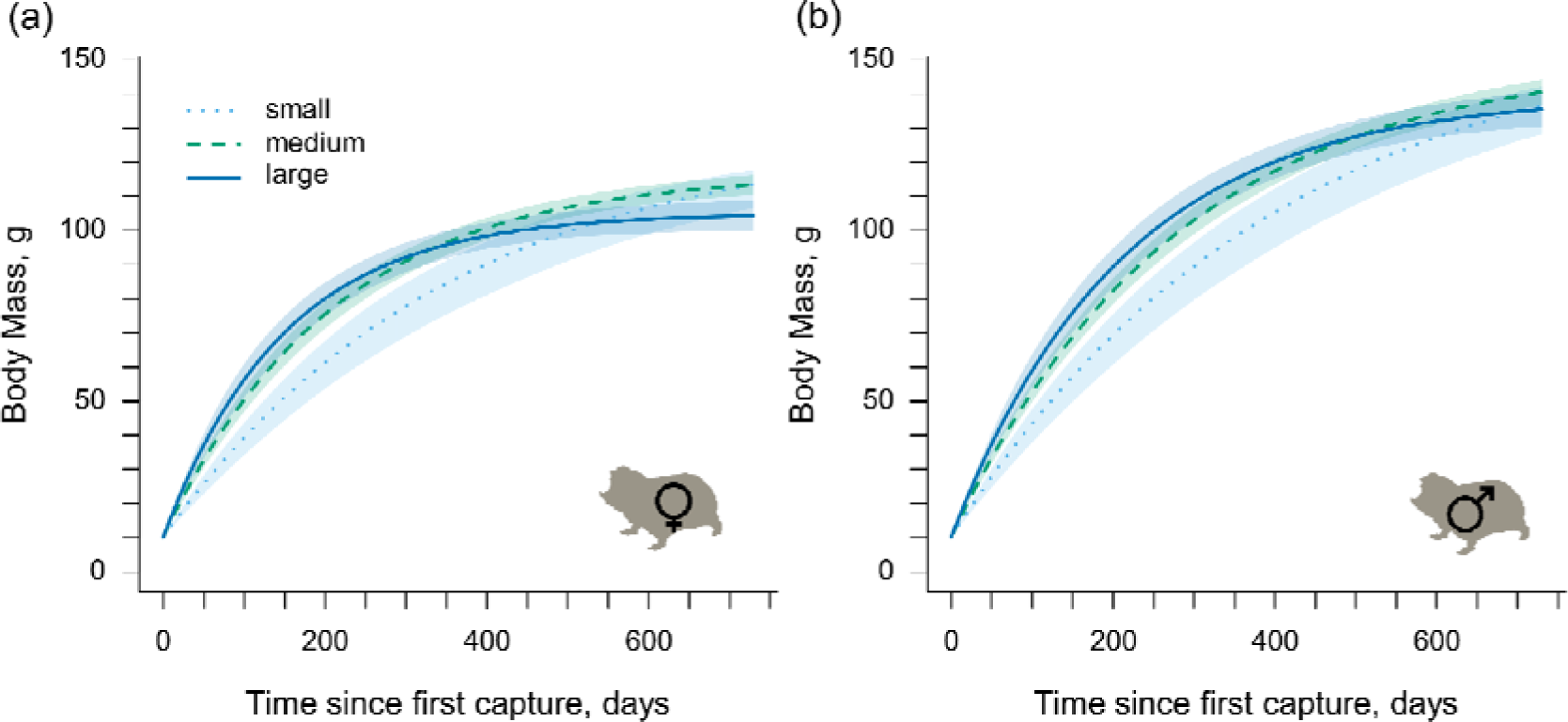
The effect on group size on body mass growth of wild Damaraland mole-rats. Body mass growth depended upon whether the individual developed in a small (4 individuals, light blue, dotted line), medium (12 individuals, turquoise, dashed line), or large (20 individuals, blue, solid line) group. (a) and (b) display the predicted body mass growth of a male or female mole-rat first captured at 10g, the average size of a pup at parturition. Curves are derived from von Bertanlanffy interval equations and show the predicted mean mass, with the shaded areas indicating 95% confidence intervals. Growth curves are estimated from models fitted to 381 mass records taken from female non-breeders in their natal group (193 unique individuals), and 456 mass records taken from male non-breeders in their natal group (214 unique individuals).

## DISCUSSION

Damaraland mole-rats have often been considered eusocial species and it has been suggested that, as in obligate cooperative breeders, successful reproduction is contingent on help provided by non-breeders (21, 35, 36, 38, 39). Our demographic analysis of wild Damaraland mole-rats shows that a substantial proportion of individuals are females that have dispersed and settled alone in a burrow system. We find that these solitary individuals display high survivorship and maintain a body condition that is comparable to individuals living in groups, despite being forced to forage alone and lacking access to other putative benefits of group-living, such as the benefits of group defence, or social thermoregulation. Once joined by a male, these nascent pairs often begin breeding immediately and produce pups at a rate which is similar to that of established groups, in spite of the lack of helpers and the known developmental costs of reproduction in inexperienced breeders (70). For this reason, Damaraland mole-rats should not be considered obligate cooperative breeders.

In demonstrating that solitary dispersal can bring reproductive rewards, our study also emphasises that female Damaraland mole-rats acquire breeding positions in ways that differ from many other cooperative mammals. In many obligate cooperative mammals, the inheritance of breeding positions - including the natal territory and pre-existing helpers - often forms the most successful route to breeding for the habitually philopatric sex, and if individuals of either sex disperse, then they typically do so in same-sex coalitions (52,53,57). Single dispersers or small coalitions have difficulty establishing new breeding groups in many of these species and the effects of group size on fitness is often initially strongly positive (3, 11, 16, 50, 71). Group founding in obligate and facultatively eusocial insects is also characterised by high chances of group failure, which are commonly reduced when several queens establish new nests together (72-74) or when colonies fission into spatially distinct breeding units (63). In our study population of Damaraland mole-rats, we found that territory inheritance was relatively uncommon in both sexes and that solitary dispersal by males and females was the most frequent route to the acquisition of a breeding position. Though solitary dispersal likely carries a high mortality risk since dispersal is often overground and mole-rats are anatomically and physiologically adapted for subterranean living, extrinsic mortality appears to be low once individuals have settled in a burrow system and have access to a supply of plant tubers.

The relative rarity of territory inheritance in our study population may be related to the low rate of successful male immigration into established groups. Most social mole-rat groups are nuclear families and females will not readily mate with their father or brothers (75). As a result, natal individuals typically lack access to unrelated mating partners and if either breeder dies, all group members may slowly disperse and groups may dissolve (21, 37, 54, 56, 61). The fact that females rarely inherit a breeding position might also explain why sex differences in dispersal timing are small in Damaraland mole-rats (this study, 68) compared to other cooperative breeders (11,57,76), and why males and females invest similarly in cooperative foraging behaviour (30,48,77), with neither sex standing to gain more from helping (11,15,76,78).

Like previous studies of social mole-rats both in captivity and in natural populations we found that rates of recruitment are higher in larger groups than in smaller ones (39) and that the work load of breeders probably declined with increasing group size (49), indicating that the presence of non-breeding helpers may contribute to breeding success or offspring survival. In addition, positive associations between group size and female fecundity have also been shown in captive Damaraland mole-rats (49). However, in many other social animals, group size, breeding success and growth often increase in groups occupying more productive ranges and habitats so that associations between group size and breeding success do not necessarily reflect causal relationships (15, 79). As yet, the extent to which the involvement of non-breeders in alloparental care contributes to the growth and survival of juveniles or to the fecundity of breeding females is unclear. However, in contrast to obligatorily eusocial insects as well as to some of the more specialised cooperatively breeding vertebrates where helpers contribute substantially to the provisioning of dependant young, this study shows that Damaraland mole-rats can survive and breed successfully without assistance from nonbreeders, suggesting that in our population the benefits of the presence of non-breeders may be small or inconsistent.

As in many other cooperative breeders (1,4,80) constraints on dispersal may play an important role in the evolution of sociality in mole-rats, as they appear to in many group-living rodents (81-83) and some carnivores (17). The high costs of dispersal may also explain why dispersal in social mole-rats is concentrated around intermittent periods of rainfall when the soil is soft enough to facilitate burrow digging (27,36,68,84), permitting the establishment of new burrows. Breeders may also tolerate non-breeders remaining in their group as a form of extended parental care, even if their retained offspring provide limited fitness benefits (7, 85).

## MATERIALS AND METHODS

### a) General methods

Damaraland mole-rats were captured in the area surrounding the Kuruman River Reserve (-26.978560, 21.832459) in the Kalahari Desert of South Africa between September 2013 and May 2020 (Fig. S1 for distribution and climate). Field work was carried out at two neighbouring localities, the Kuruman River Reserve (“Kuruman”), and the Lonely farm (“Lonely”, Fig. S2 & S3). The two locations form a continuous population but because of the time intensive nature of mole-rat trapping they tended to be trapped in discrete periods (Fig. S4 & S5), hereafter referred to as “trapping windows”. For logistical reasons, trapping stopped at Lonely in July 2017.

Groups of mole-rats in the wild are revealed by the lines of mounds that they extrude when excavating their tunnel systems (Fig. 1). Groups were trapped periodically at 6 to 12-month intervals using modified Hickman traps that were baited with sweet potato and positioned into tunnel systems once accessed by digging. After trap setting, traps were checked every 2-3 hours throughout the day and night. On capture, animals were placed into a closed box and provided with food and shelter. Intermittently, individuals were transported back to the laboratory where they were sexed, weighed to the nearest gram and measured for various morphometrics. Breeding females are commonly the largest female in their group and can be easily identified from their perforated vagina and prominent teats. All captured individuals were marked with passive integrated transponder (PIT) tags on first capture to allow for individual recognition on recapture. After sampling, groups were housed temporarily in semi-natural tunnel systems in the laboratory and provided with food, nesting material and sand. The complete group was assumed to have been captured after an absence of activity at the trap sites for 24 hours, after which point the animals were returned to their tunnel system. The average time to capture a group was 2.73 ± 1.67 days (mean ± SD).

Between September 2013 and May 2020, 752 unique individuals (368 females, 384 males) were trapped at the two sites (N = 1941 individual captures). The mean recapture rate of individuals across successive trapping windows was 73.1 ± 10.8% at Kuruman and 54.8 ± 14.4% at Lonely (mean ± SD; Table S1). In total, analyses used data from 328 group captures that were carried out at 84 groups with an average of 3.90 ± 2.99 captures per group (mean ± SD; Fig. S2). Groups were defined as individuals repeatedly found at the same trapping location, and in the majority of recapture events the same breeding female remained at a group for their duration. Thus, as long as the majority of individuals trapped within the same area were known to originate from the same group, the group continued to be defined as such. On several occasions a new group had moved into the tunnel system previously occupied by another group. The initial group were then either found elsewhere or never recaptured and likely extinct. Single individuals, either new unknown individuals or known individuals that had dispersed from an established group were assigned their own unique group ID, which was retained if they were later joined by an immigrant partner and started breeding in the same area.

In 15.2% of group captures, continued activity at trapping sites indicated that the complete group had not been captured by the end of the week of trapping. The decision to include or exclude these incomplete captures depended on the analysis: group-level analyses excluded incomplete captures whereas individual-level analyses included them (a summary of the data and models used for the various analyses is provided in Table S2.). Analyses were conducted in R version 3.6.3 (86), and all code is provided online (https://github.com/JThor1990/DMR_GroupSizeEffects). Summary statistics of the raw data are reported as the mean ± SD; model estimates are reported as either the mean ± 1SEM, or as a hazard ratio ± 95% confidence intervals.

### b) Status-related survivorship among females

To estimate the survivorship of females in different states we fitted a multi-state Markov model (MSM). Such models are typically used in a medical setting for ‘panel data’, where a continuously observed process - like disease progression - is measured only at discrete time points - when people choose to visit the hospital. The timing of transitions between states can then be estimated indirectly under the assumption that the next state in a sequence of states depends only on the current state, and not on the history of transitions or on the time spent in the prior state (87). Our longitudinal data bears a panel-like structure, with individuals and groups being periodically captured (or not) in different life history stages or ‘states’. The model then estimates the probability that individual mole-rats transition between different states. Here, we present information on survival probability from each state.

Information on individual capture histories was used to assign females to one of four states relative to the time since their first capture: *i*- *non-breeder* in their natal group, ii- *single female*, *iii*-*breeding female* out of their natal group, *iv*-*disappeared*/*dead* (Fig. S5). For model fitting purposes, in the small number of cases where females inherited a breeding position within their natal group (n = 7), these females were categorised in the same state as females that acquired a breeding position out of their natal group (*iii*). To define state *iv* (disappeared/died), we incorporated information from the trapping windows at “Kuruman” and “Lonely”. If an individual had not been recaptured for at least two consecutive trapping windows, it was assumed to have dispersed or died at some point between its last live capture and the start of the subsequent trapping window, and given an extra row of data reflecting this; the model is then parameterised so that the timing of disappearance or death is not exact. Similarly, if individuals were captured within the last trapping window at each location, we assumed that they were still alive at the end of that trapping window (i.e. the end of the study). They were then given an extra row of data reflecting this assumption and were censored at this point, which in the most extreme case assumes that individuals remained in their last captured state for a further 80 days (Fig. S6). The multi-state analysis included 326 females that were captured as a *non-breeder* in their natal group at some point in the study, 45 females captured as a *single female*, and 46 captured as a *breeding female*. To explore whether group size affected the likelihood that non-breeding females or breeding females died/disappeared, we fitted an additional model that included a group size covariate for each of these transitions. The multi-state models were fitted in the *msm* package (87; Fig. S7 for multi-state diagram). When reporting hazard ratios and relative likelihoods from the multi-state models we class cases where the 95% confidence intervals do not overlap one as indicating a biologically important effect, while also emphasizing effect sizes. To directly compare survivorship among the different classes of female, we also calculated the ratio of the transition intensities from each alive state (*i*, *ii*, or *iii*) to death/disappearance (*iv*), with 95% confidence intervals computed using the delta method (*qratio.msm* function).

### c) Within-group recruitment

Two approaches were used to investigate the role of group size on within-group recruitment. In the first, recruitment was analysed longitudinally across the duration of the study. In the second, new groups were experimentally created in the wild through the introduction of unfamiliar adult males to solitary females. The recruitment rate of newly created pairs was then compared to that of established groups that were captured and recaptured within the same time period. The benefit of the second approach is that it allows for a direct test of whether groups in the early stages of group formation were less productive than established groups. Any individuals caught within one year that were lighter than 100g (males) or 80g (females) were assumed to be recruited from within the group, as informed by the growth curves. By contrast, any new individuals heavier than these cut-offs were assumed to represent out-of-group immigrants.

The longitudinal analysis used information from every capture-recapture event where a group was captured and then recaptured within a 100 to 365-day time period; where a resident breeding female was present; and where at least one large male was retained across the two captures. The analysis is therefore restricted to active breeding units. In total, this produced a dataset of 78 group-level capture-recapture events that took place in 33 groups with a mean trapping interval of 214.2 ± 52.1 days (mean ± SD). The number of recruits was modelled using a generalised linear mixed effects model (GLMM) with Poisson error in the *glmmTMB* package (88). Fixed effects of group size at first capture, the mass of the breeding female at the capture, and rainfall were included. A single random effect of group identity was included, and the logged time interval between capture and recapture was included as an offset term. Rainfall was calculated as the geometric mean monthly rainfall in the year preceding the first capture as this provided the best fit to the data (lowest AIC score) when compared to models fitted to the total rainfall, or the arithmetic mean rainfall. After standardising relative to the trapping duration, the mean 6-monthly recruitment to groups was 2.12 ± 1.82 individuals (mean ± SD). Continuous variables were z-score transformed prior to model fitting.

For the experimental approach, 8 solitary females were captured between 16^th^ April 2016 and 29^th^ May 2016. Solitary females were brought back into the laboratory and paired with wild males captured concurrently from intact breeding groups. When pairing, males and females were first isolated in their own temporary tunnel system and exposed to the odour of their prospective partner for 24 hours. After this time period, the male was introduced to the female for 24 hours in her lab tunnel system, before they were returned together to the female’s tunnel system in the wild. All pairings used large adult males originating from groups > 3km from the focal solitary female, so it is highly unlikely that pairings were conducted between close relatives and/or familiar individuals. The recruitment rate of new pairs was compared to that of established groups captured over the same time period using a Welch’s t-test after first standardising recruitment rate to a yearly measure. Groups where a male was removed were not included in the established groups treatment in case the male was a breeder in his original group, which would necessarily reduce the rate of recruitment. The mean trapping interval did not differ between new pairs (249.89 ± 94.34 days; mean ± SD) and established groups (306.88 ± 64.39 days; mean ± SD; Welch’s t-test: t = -1.47, df = 14.14, p = 0.16).

### d) Early-life growth

As the age of wild-caught individuals was unknown, we modelled growth using interval equations that estimated the change in body mass and upper incisor width of individuals across successive capture events. The upper incisors are a reliable measure of skeletal size in Damaraland mole-rats and are the main apparatus of digging (89). Incisor width was measured at the widest point using digital callipers (to the nearest 0.01mm). All measurements were taken in duplicate by two observers and the mean of these two measurements was used. Previous studies show that the shape of growth in captive mole-rats is concave and can be approximated by a monomolecular curve (90). To allow for a similar shape of growth in the wild, the interval equation was parameterised as a von Bertalanffy growth curve (91) and fitted as a non-linear mixed effect model (NLMM) in the *nlme* package (92). The body mass or incisor width of an individual at each recapture was estimated as:

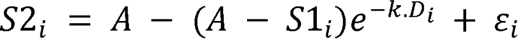

where S1*_i_* and S2*_i_* are the size at capture and recapture, respectively, for individual *i*. A is the estimated population-level asymptotic size, *k* is the growth rate constant, *D_i_* is the time difference between capture and recapture (hereafter the ‘trapping interval’). ε_*i*_ is the normally distributed error with mean 0 and variance σ^2^.

Separate models were fitted to males and females for each size metric, using all information from individuals that were recaptured at least once, and where the recapture interval fell between 90 and 365 days. The female body mass dataset comprised 381 “repeat capture” events (*n* = 193 females, mean/female = 1.96 ± 1.21; mean ± SD) and excluded any weight data from females once they were known to be a breeder, thus removing any artefact of status or pregnancy. The male body mass dataset consisted of *n* = 456 repeat capture events (*n* = 214 males, mean/male = 2.13 ± 1.54; mean ± SD). The female and male incisor width datasets consisted of 328 and 381 repeat capture events respectively (n = 193 females, mean/female = 1.82 ± 1.16; n = 198 males, mean/male = 1.92 ± 1.33; mean ± SD). In each model a random effect of individual identity was specified at the level of the asymptote and the growth rate constant, and to aid convergence, random effects were modelled as uncorrelated. Likelihood ratio tests indicated that the inclusion of a random effect of group identity was not needed.

The initial growth model was extended by incorporating a standardised group size term:

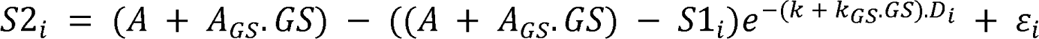

Here, *A*_GS_ and *k*_GS_ estimate the change in asymptotic mass and growth rate constant with changing group size, *GS*. Group size was taken as the group size on initial capture, which was highly correlated with recapture group size (r = 0.78, df = 835, p < 0.001). As the results were qualitatively unaffected by excluding incomplete captures, results are presented for all captures. Group size was z-score transformed prior to model fitting, such that *A* and *k* are estimated at the mean group size experienced by each sex.

## Acknowledgements

We are grateful to numerous field assistants that helped to collect the field data (usually under hot and/or nocturnal conditions), and to numerous research managers and onsite employees who have contributed to the long-term running of the Kalahari mole-rat project. In particular, we would like to acknowledge Philippe Vullioud, David Gaynor, Tim Vink, Walter Jubber, and Nigel Bennett for various logistical and academic input to the project. We are also indebted to the Kalahari Research Trust for access to the facilities, and to Prof. Marta Manser for her contributions and efforts towards running and maintaining the Kalahari Research Centre, supported by funding from the University of Zurich and the MAVA foundation. Lastly, we thank the Northern Cape Department of Environment and Nature Conservation for permission to work in the Northern Cape, and Kobus Lamprechts for allowing us to work on his land. This work was supported by grants from Vetenskapsrådet (2017-05296) and Crafoordska Stiftelsen (2018-2259 & 2020-0976) awarded to M.Z., and grants from the European Research Council (European Union’s Horizon 2020 research and innovation program, no. 742808 and no. 294494) awarded to T.C-B.

## ETHICS

The research carried out in this study was approved by the University of Pretoria Animal Ethics Committee (ECO32-13, EC050-16)

## Supporting Information

**Fig. S1.**
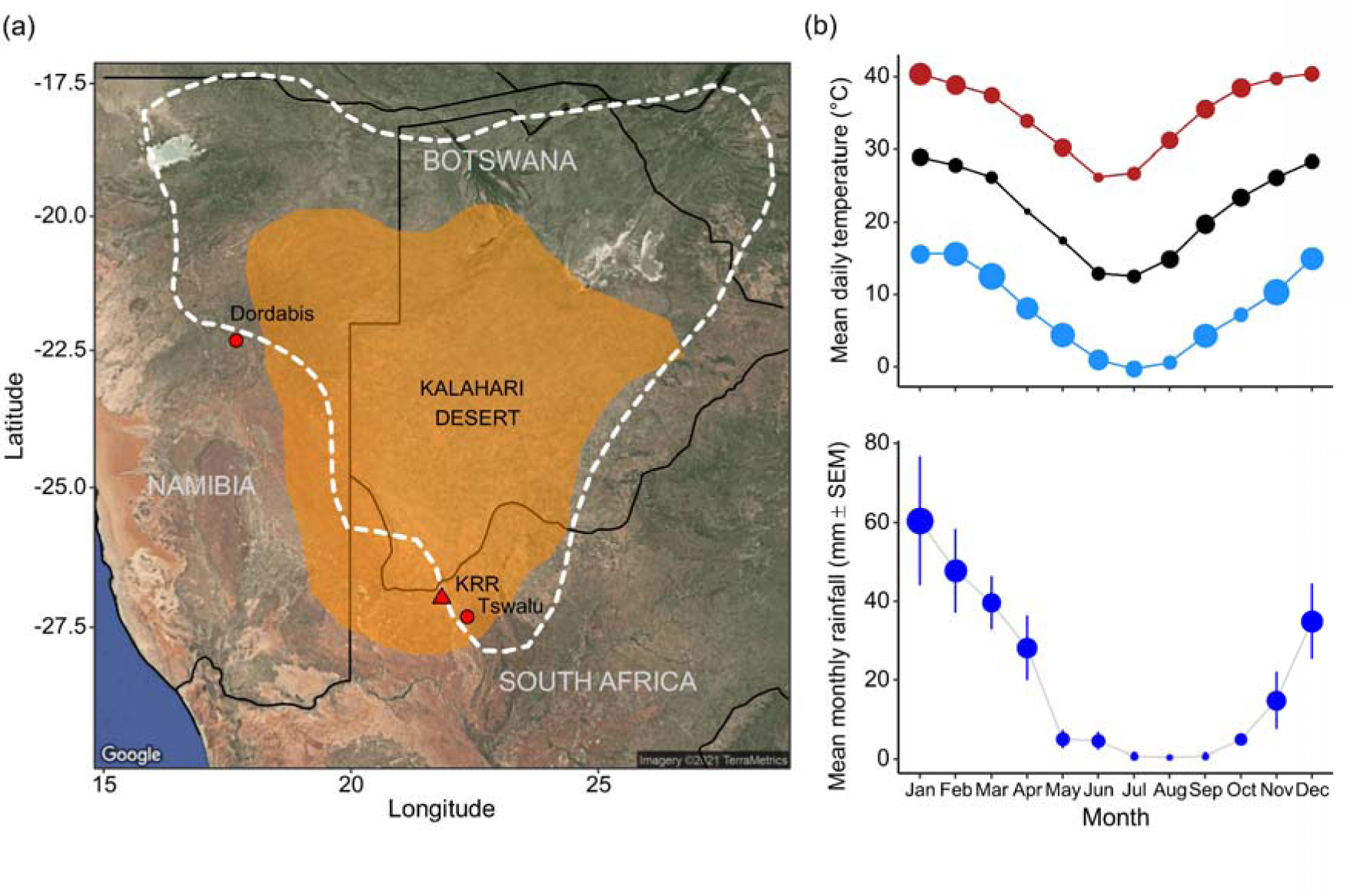
a) The distribution of the Damaraland mole-rat in sub-Saharan Africa. The predicted current distribution is highlighted by the white dashed line [taken from IUCN, 30^th^ June 2021], with the Kalahari Desert indicated by the shaded polygon. The location of the Kuruman River Reserve (KRR) is also highlighted, as are two other sites where medium to long-term capture-mark-recapture studies of Damaraland mole-rats have been carried out. b) The climate at the KRR between 2010 and 2020. Upper panel shows the mean, minimum, maximum daily temperature in each month, averaged across the years, and the lower panel shows the total monthly rainfall (± SEM), averaged across years; points are scaled to 1SEM. Climate data taken from NASA’s GMAO MERRA-2 assimilation model and GEOS 5.12.4 FP-IT.

**Fig. S2.**
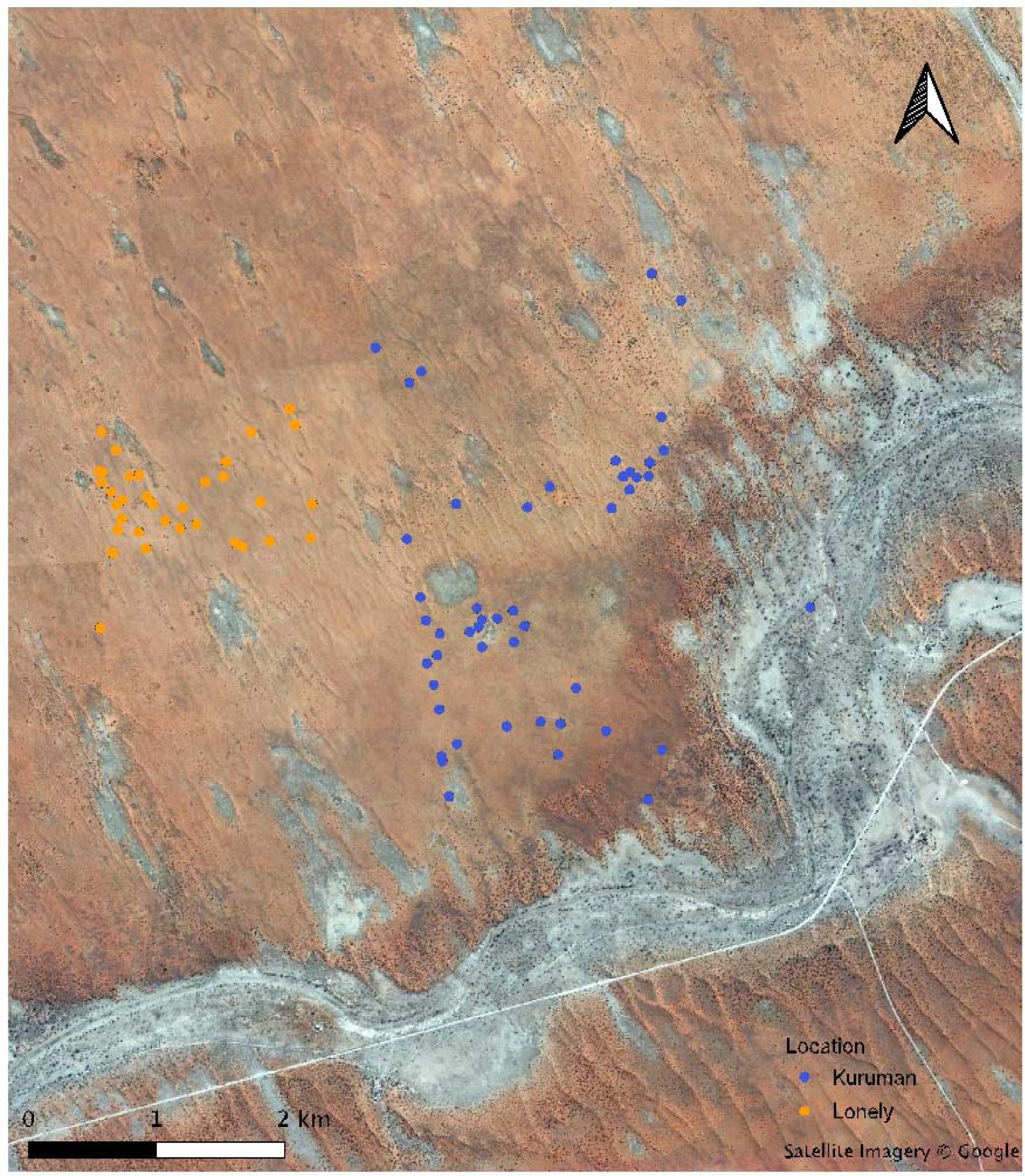
The trapping locations of Damaraland mole-rats groups captured at the Kuruman River Reserve and Lonely farm between 2013 and 2020. Each group is noted by a point which marks the centroid of all group captures, with the point colour corresponding to location, “Kuruman” or “Lonely”. Note that with only one exception, mole-rat groups were distributed entirely within the red ‘arenosol’ soils, where their principal food (Gemsbok cucumber) is found. The one exception to this pattern was a single dispersing female who temporarily burrowed in an area of calcareous soils, which appear light grey from satellite. These calcareous soils highlight the path of the long dried up Kuruman river as it passes through the reserve.

**Fig. S3.**
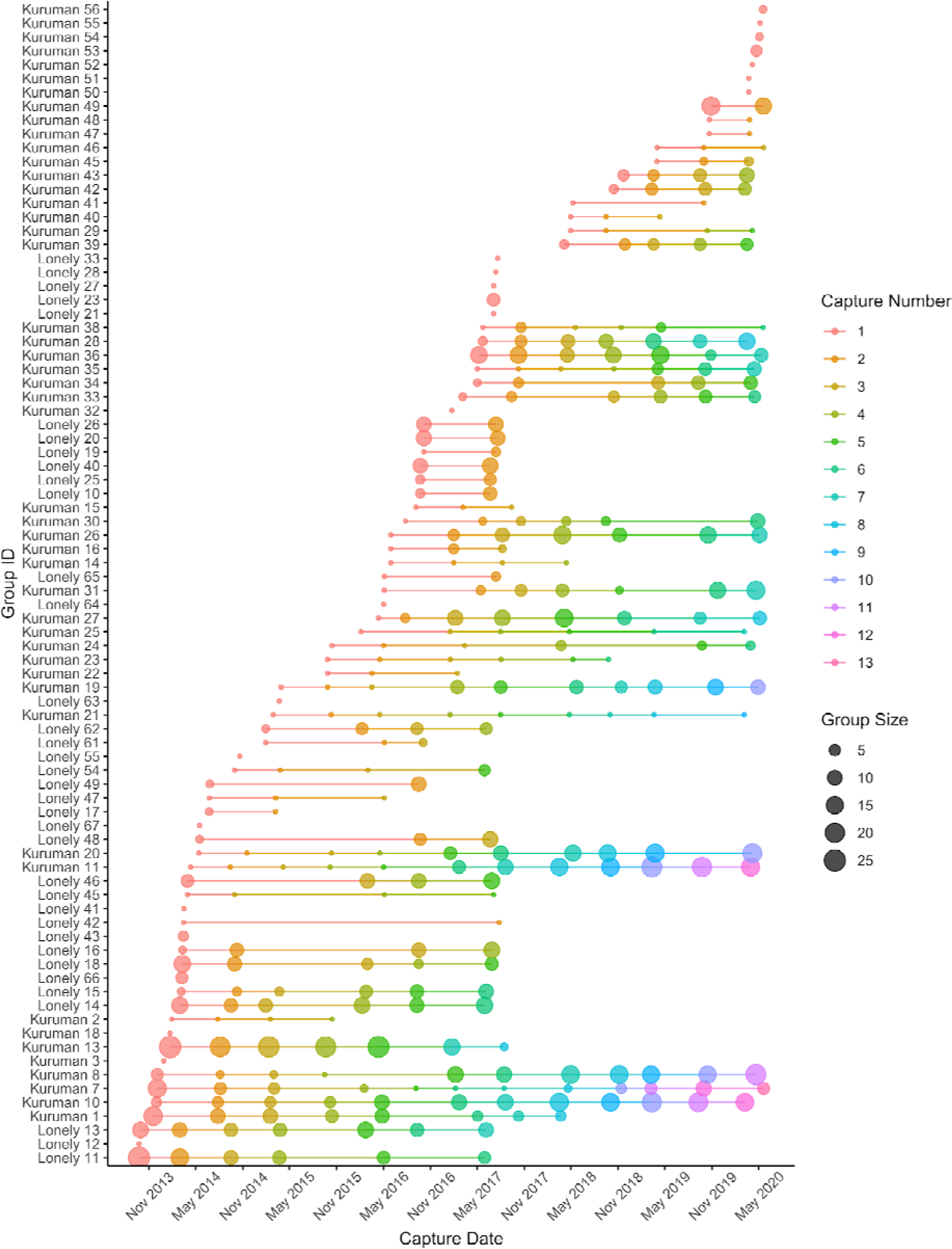
The history of trapping effort of Damaraland mole-rats groups in and around the Kuruman River Reserve. Each unique group has its own row, where successive capture events are denoted by points moving right along the x-axis. The size of the points is proportional to the size of the group at each capture event.

**Fig. S4.**
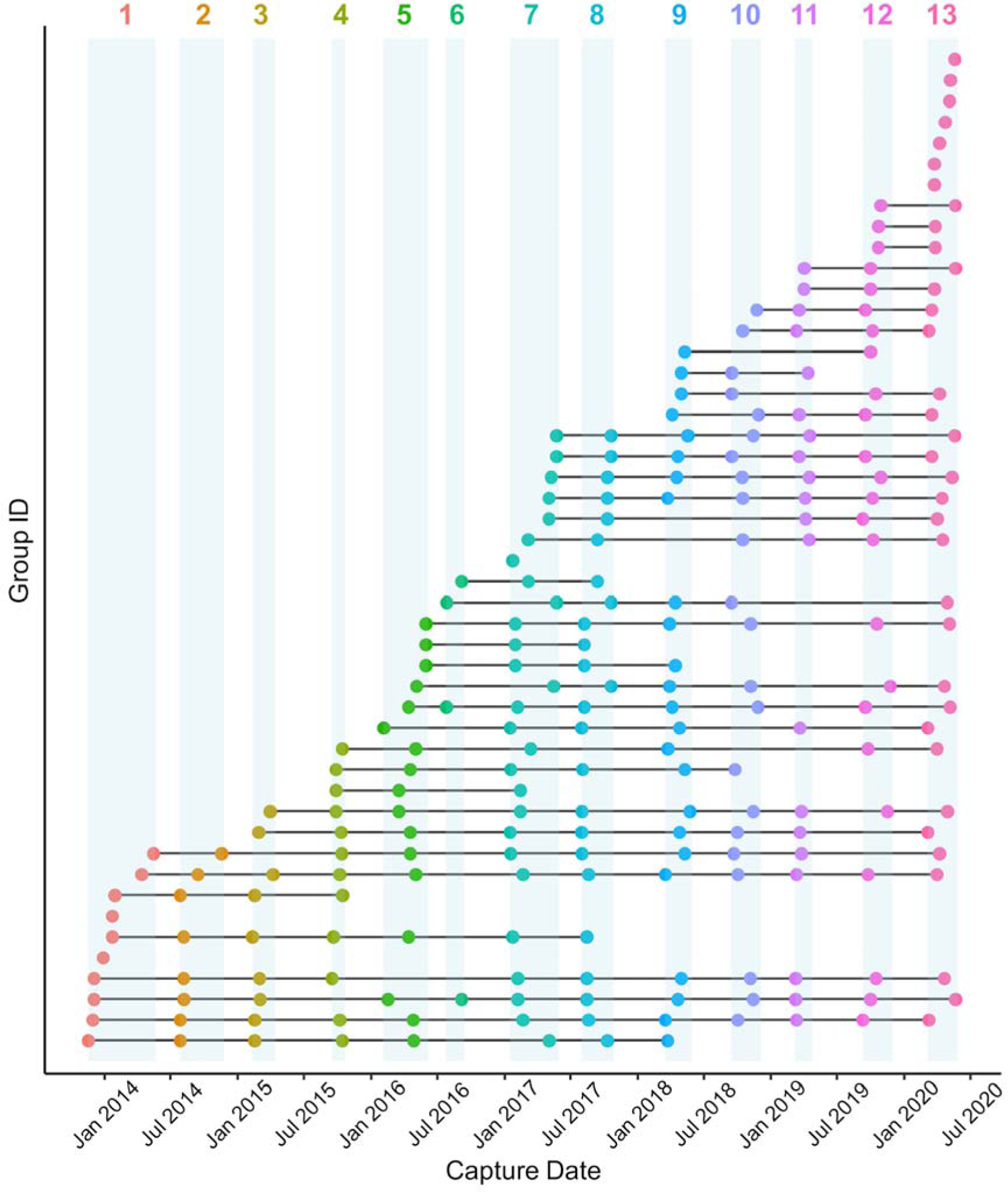
The history of trapping effort for Damaraland mole-rat groups at the Kuruman River Reserve in the Kalahari, separated into discrete trapping windows. Periods are defined by any break in trapping effort of more than 2 months. By doing so, it can be seen that no group is captured twice within a trapping period. Also, most groups are captured in successive trapping periods, with very few groups ‘skipping’ trapping windows-with the exception of period 6, where only a few groups were recaptured. In the few other cases where groups skip a trapping period, this is because no mounds were visible throughout the period, and when this occurs, it is not known whether a group had collapsed/disappeared or was present underground but not digging.

**Fig. S5.**
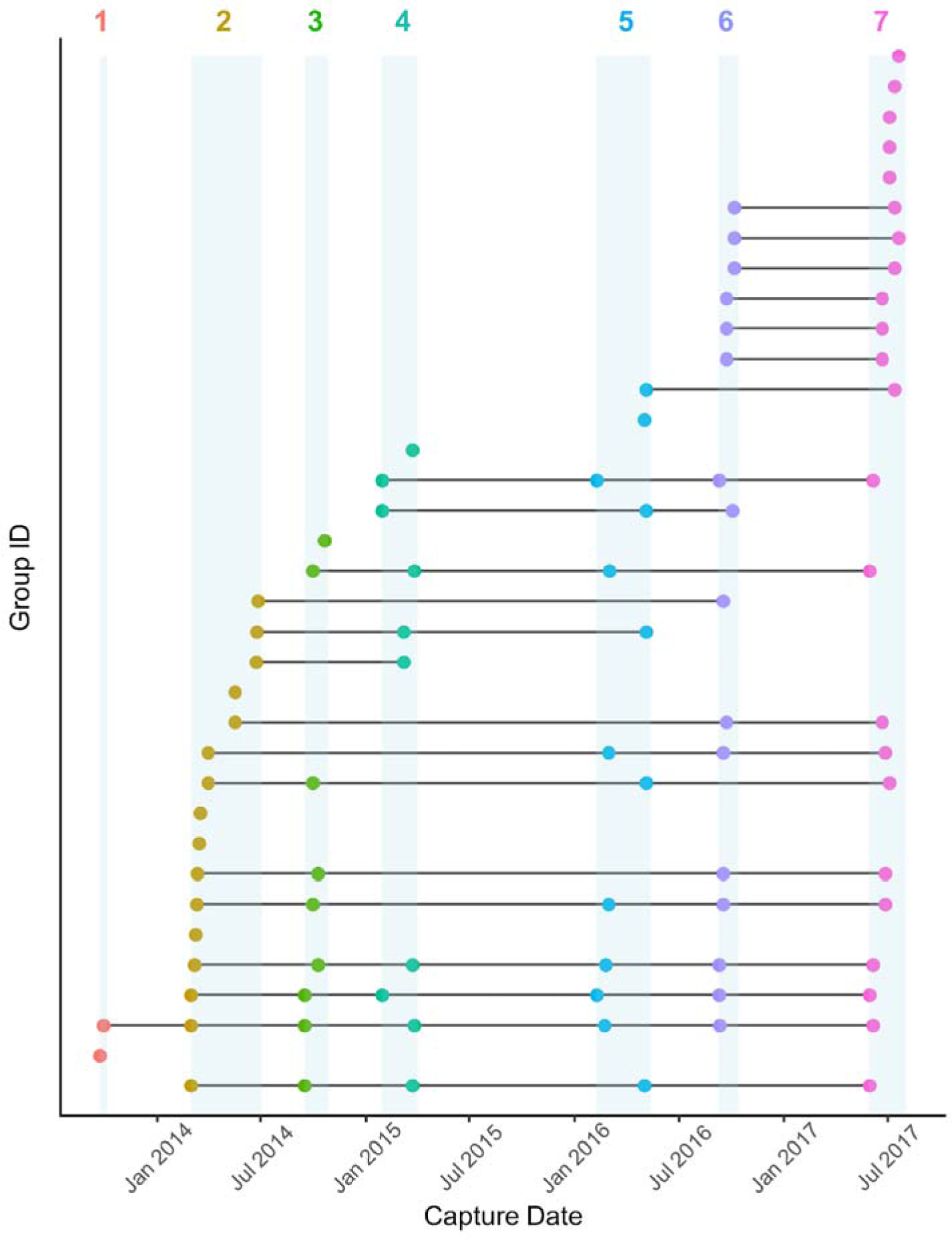
The history of trapping effort for Damaraland mole-rat groups at “Lonely” in the Kalahari, separated into discrete trapping windows. Periods are defined by any break in trapping effort of more than 2 months. By doing so, it can be seen that no group is captured twice within a trapping period. Also, most groups are captured in successive trapping periods, with very few groups ‘skipping’ trapping windows-with the exception of period 6, where only a few groups were recaptured. In the few other cases where groups skip a trapping period, this is because no mounds were visible throughout the period, and when this occurs, it is not known whether a group had collapsed/disappeared or was present underground but not digging.

**Fig. S6.**
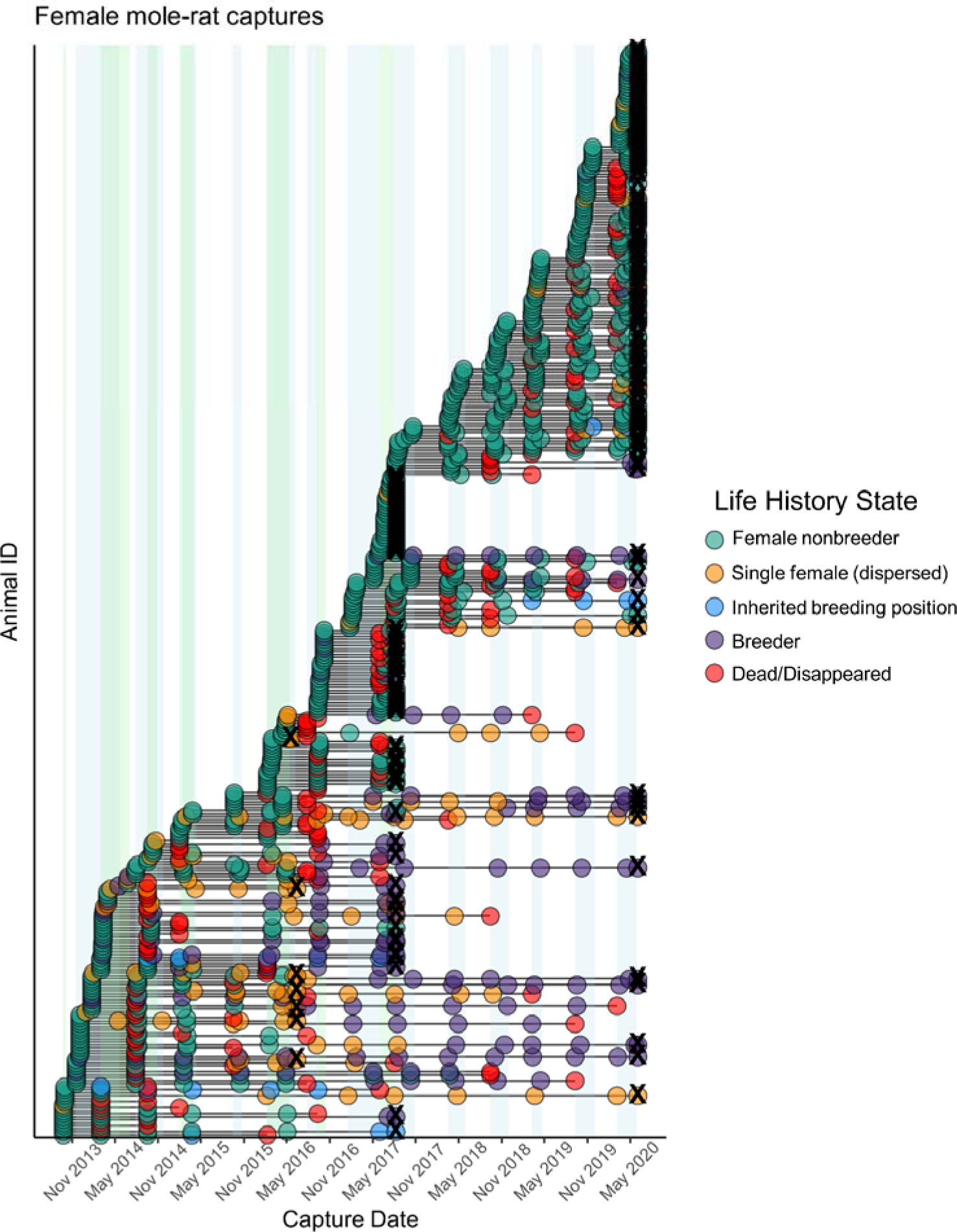
The fates of female Damaraland mole-rats capture from 2013-2020. Though analysed together, the fate/current state of female non-breeders at the Kuruman River Reserve and Lonely were assessed separately since they took advantage of different trapping windows (KRR-blue shaded areas; Lonely; green shaded areas). Individual states at any given time are female non-breeder, single female (who has dispersed from her natal group), out-of-natal group breeder (again dispersed), and disappeared/died. In a small number of cases females also acquired dominance in their natal group (when an immigrant male had moved in previously). For individuals that were not recaptured at some point before the final trapping period at either location, it was assumed that the individual had died or disappeared at some point between their last capture and start of the next trapping window; as handled by the models. Additionally, individuals captured in the last trapping period at each location were assumed to have persisted in their current state up until the end of the period, where they were then right-censored (**X**).

### Multi-state modelling

In the Materials and Methods in the main text we outline the fitting of a multi-state model to investigate the survivorship of females in different classes or life history stages (Fig. S6). This model relied on the assignment of females as either (i) an in-group non-breeder, (ii) a dispersed single individual, (iii) a breeder, or (iv) died or dispersed-using information on capture histories (Fig. S5).

**Fig. S7.**
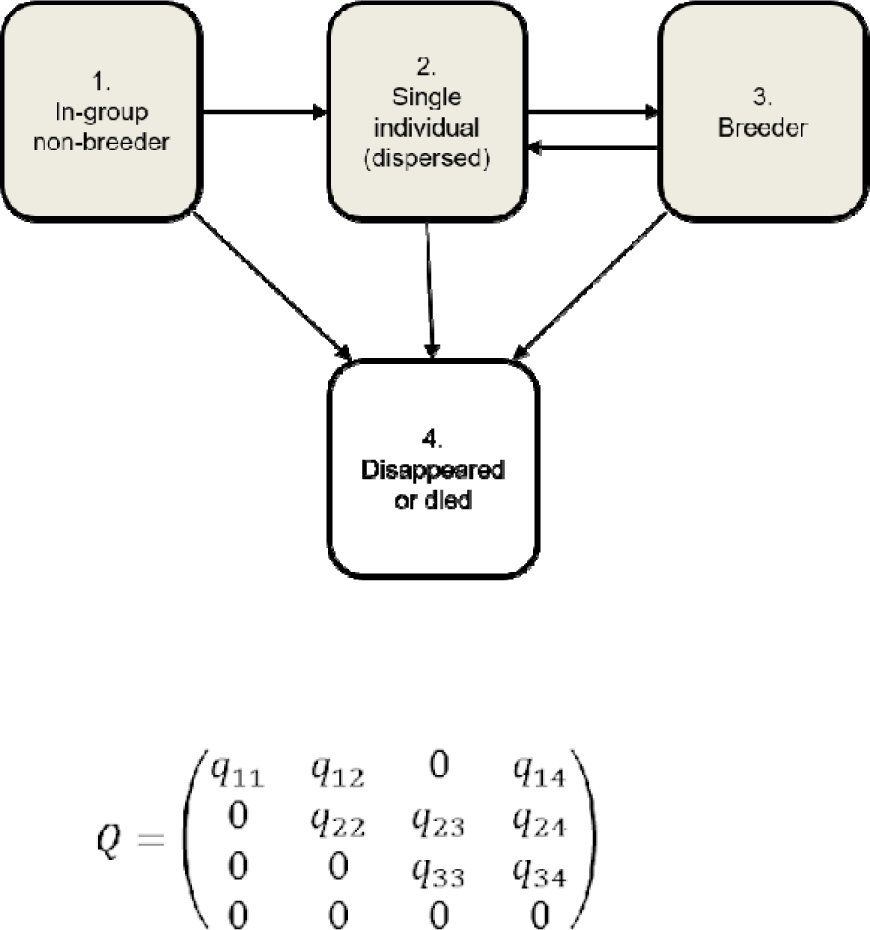
Multi-state model for the life history transitions of female Damaraland mole-rats, with associated Q matrix. In a small number of cases where individuals were experimentally paired (i.e. given a male when formerly present as a single female and analysed in other papers; e.g. Thorley et al. 2018 Proc. R. Soc. B, 285: 20180897), they were also right-censored at the point of pairing so that any transitions from single female to breeder did not include this experimental component. In all cases, the multi-state model was parameterised so that the timing between transitions was not known exactly, but was estimated under Markov assumptions-this includes the timing of death/disappearance.

### Dispersal timing

We adopted a similar state-based framework to investigate sex differences in the timing of natal dispersal. In order that the timing of dispersal could be approximately related to age, these analyses focussed on individuals that were first captured in their group at less than one year of age (<80g for females, <100g for male; Fig. 5 main text). Unlike the female survivorship model, where individuals were assigned to one of two states, the philopatry model defined only two states for males and females: i-*non-breeder* in their natal group, and ii-*disappeared*/*dead/known disperser*. Any conditions under which an individual first left their natal group or disappeared was specified as state (ii). Males and females were modelled together, and a covariate for sex was included so that the transition intensity of males was estimated relative to females. Covariates of weight and rainfall were also included to infer whether the disappearance of non-breeders from groups was mostly due to dispersal or to death in situ; if dispersal was the principle reason for disappearance, then we expected that heavier (and older) individuals would be more likely to disappear [or be successfully recaptured out of their natal group]. We also predicted that increases in rainfall were associated with higher rates of disappearance, which we reasoned would also be suggestive of dispersal rather than death in situ. Rainfall was characterised as the total rainfall (mm) which fell three months prior to leaving state (i).

The dispersal model indicated that males and females disappeared from their natal group at a similar age (Fig. S7). For individuals first captured at less than one year of age, 39.5% of females were predicted to have disappeared one year later (95% CI = [33.5,45.5]), as compared to 43.5% of males (95% CI = [38.3,49.3]). This sex bias reflected a tendency for males to leave their natal group slightly earlier than females, though this effect was not statistically significant (MSM: hazard ratio of males relative to females = 1.125, 95% CI = [0.864,1.476]; Table S1).

**Table S1.**
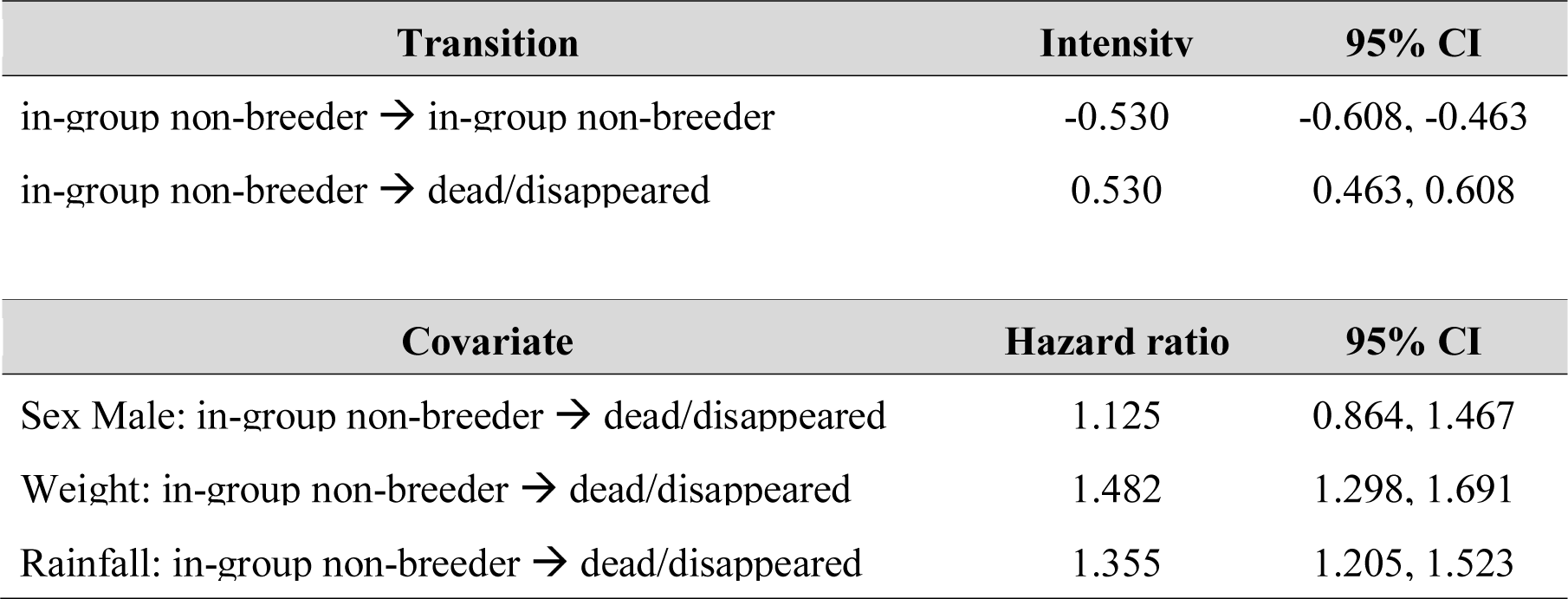
Duration of philopatry in males and females. Incorporation of weight and rainfall terms. Transition intensities from a ‘multi-state’ model fitted to non-breeders first captured at less than one year of age (here only two states considered). Prior to model fitting, the weight term was standardised within each sex, and rainfall was standardised across both sexes. Rainfall represents the total (sum) daily rainfall in the 3 months prior to disappearance/dispersal.

**Fig. S8.**
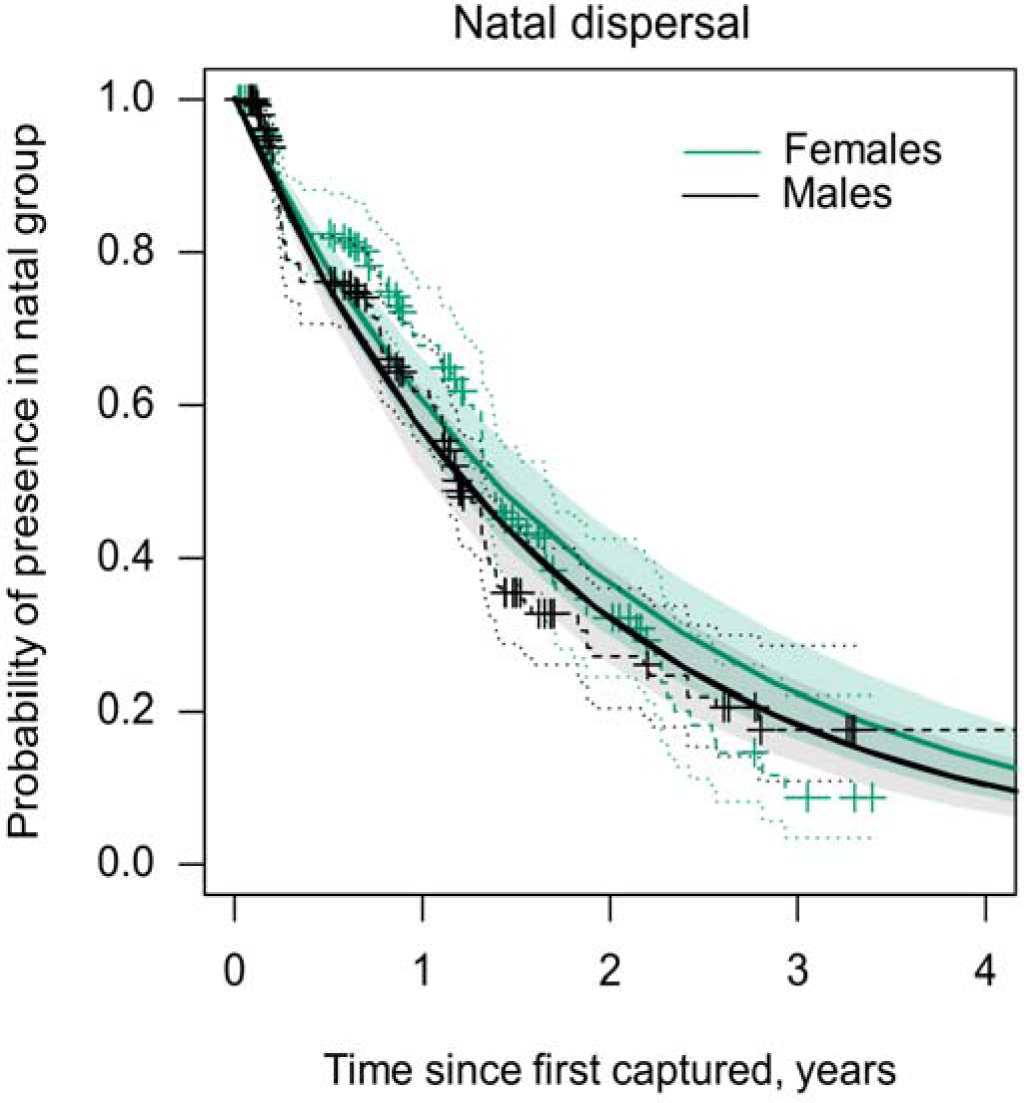
The duration of philopatry (timing of dispersal) for female (green) and male (black) Damaraland mole-rats at the Kuruman River Reserve. Solid lines display the expected probability of recapturing non-breeding individuals within their natal group, as estimated from a Markov model fitted to all individuals first captured at less than one year of age (females <80g, males < 100g). Dotted lines display the ‘empirical’ Kaplain-Meier estimate of survival probability, with crosses denoting cases of censorship. Disappearance combines cases of in-group mortality with cases of dispersal, where individuals have left their current group and state and are either re-captured subsequently in a different state, or not re-captured thereafter.

The mean timing of dispersal for males and females that is presented in the main text was estimated as the mean sojourn time in the natal group (*sojourn.msm()* function), when rainfall and body mass were held at their mean value across the data set. For males, the mean sojourn time in the natal group was 1.80 years (95% CI [1.50,2.15]) after their first capture, and for females was 2.02 years (95% CI [1.66, 2.47]). With groups being trapped at roughly 6 month intervals, these values reflect an underestimate of the average age at dispersal.

The incorporation of weight and rainfall terms into the multi-state model suggested that a large proportion of the non-breeders that disappeared from groups were individuals that dispersed, rather than individuals that died in situ (Table S1). Firstly, heavier individuals were more likely to disappear from their natal group than lighter individuals (MSM: hazard ratio = 1.482, 95% CI [1.298,1.691]), as would be expected if dispersal was more prevalent in older (heavier) individuals. Secondly, individuals were more likely to disappear following periods of increased rainfall (MSM: hazard ratio = 1.355, 95% CI [1.205,1.523]).

### The condition of single females

To compare the body condition of single females and in-group non-breeding females we performed standardised major axis regressions of body mass on two skeletal size traits: incisor width and body length. We included body length as this is a more conventional linear measurement of skeletal size in mammals. Incisor width was measured as noted in the main text. Body length was measured dorsally from the front of the snout to the tip of the tail using a tape measure (to an accuracy of 1mm). As for incisor width, each measurement was taken in duplicate by two observers, and we here use the average of these two measures. For each skeletal measurement we filtered out any in-group non-breeding females whose skeletal size was less than the minimum size trait found in single females. This meant that the two classes of female were size-matched skeletally, such that any difference in condition would reflect increased bodily reserves for a given skeletal size.

Standardised major axis regression was proposed by Peig & Green (2008) as a superior regression method for the scaling of mass on linear body measurements than ordinary least squares methods, and has been widely adopted in studies on mammals since. Briefly, we performed a standardised major axis regression of ln body mass against the natural log of each skeletal measurement using the *smatr* package (Warton et al. 2012). In both cases, we initially fitted a model which allowed the slope and intercept to vary according the class of female, but for both skeletal measurement neither the intercept (teeth width LRT: χ^2^_1_ = 1.04, p = 0.31; body length LRT: χ^2^_1_ = 0.09, p = 0.76) nor the slope (teeth width LRT: χ^2^_1_ = 1.31, p = 0.25; body length width LRT: χ_21_ = 2.53, p = 0.11) significantly different between single females and in-group non-breeders.

### This indicates that single females and in-group non-breeders did not differ in their condition

We therefore present the results of the scaling relationship where all females were assumed to following the same scaling relationship; modelled by a single slope and intercept (Fig. 3 in main manuscript, incisor width intercept = 2.41, 95% CI = -0.08 – 0.56, incisor width slope = 2.57, 95% CI = 2.39 – 2.75; body length intercept = -4.83, 95% CI = -5.33 – -

4.33, body length slope = 3.34, 95% CI = 3.17 – 3.52).

The scaled mass index of each measurement can be calculated using the following formula:

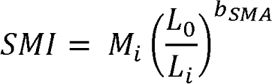

Where *M*_i_ and *L*_i_ are the body mass and skeletal measurement of individual *i*, *b_SMA_* is the scaling exponent estimated by the SMA regression of *M* on *L*, and *L*_0_ is the arithmetic mean value of the population sample. The scaled mass index of single females and in-group non-breeding females are contrasted below in the inset boxplots.

### Experimental Pairings

**Fig. S9.**
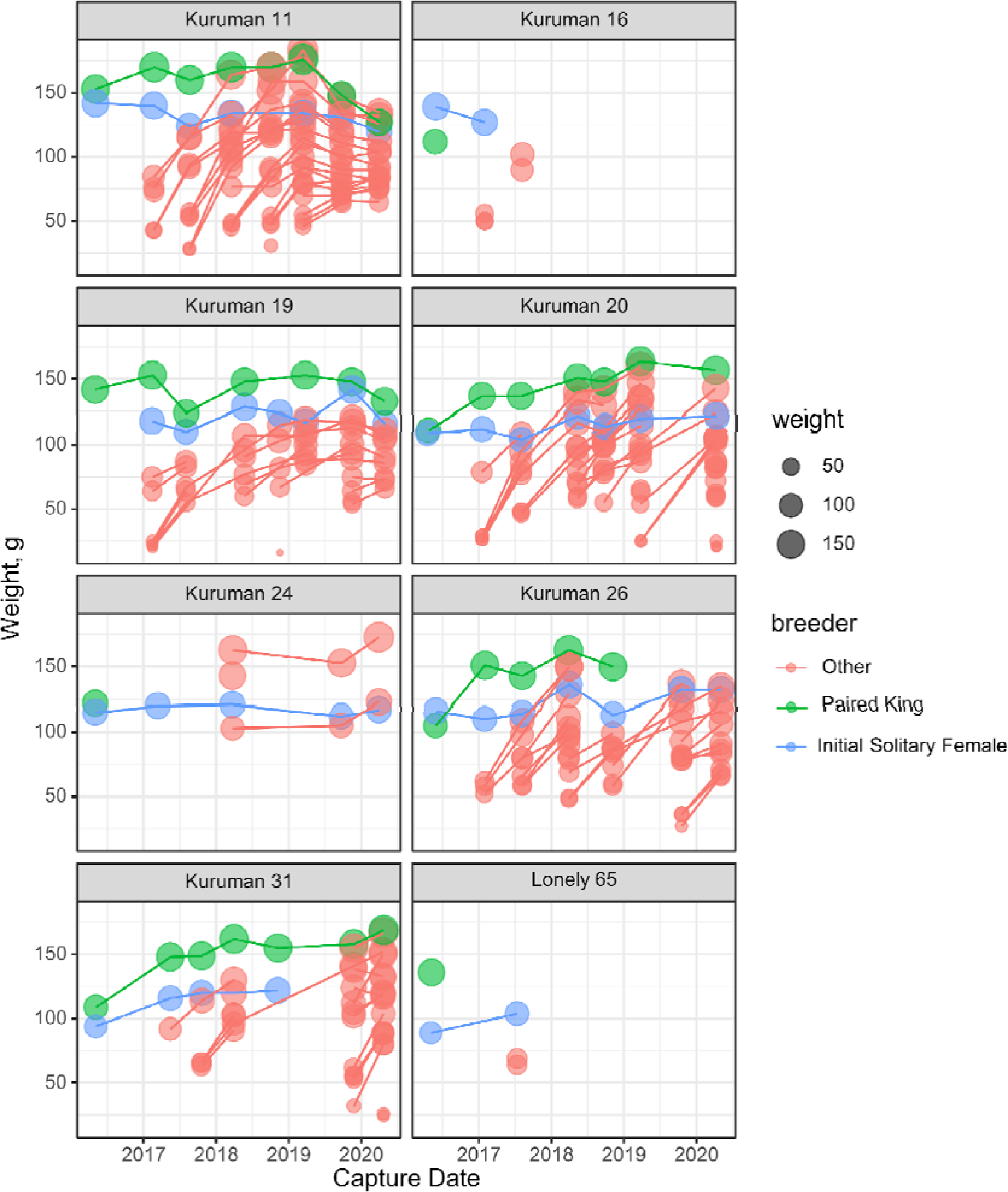
Group structure and trapping history of groups that were created experimentally with the addition of an unfamiliar male to the burrow systems of single females. In most cases newly created pairings commenced breeding shortly after pairing, as indicated by the recruitment of ‘other’ individuals into the group following the initial pairing.

**Fig. S10.**
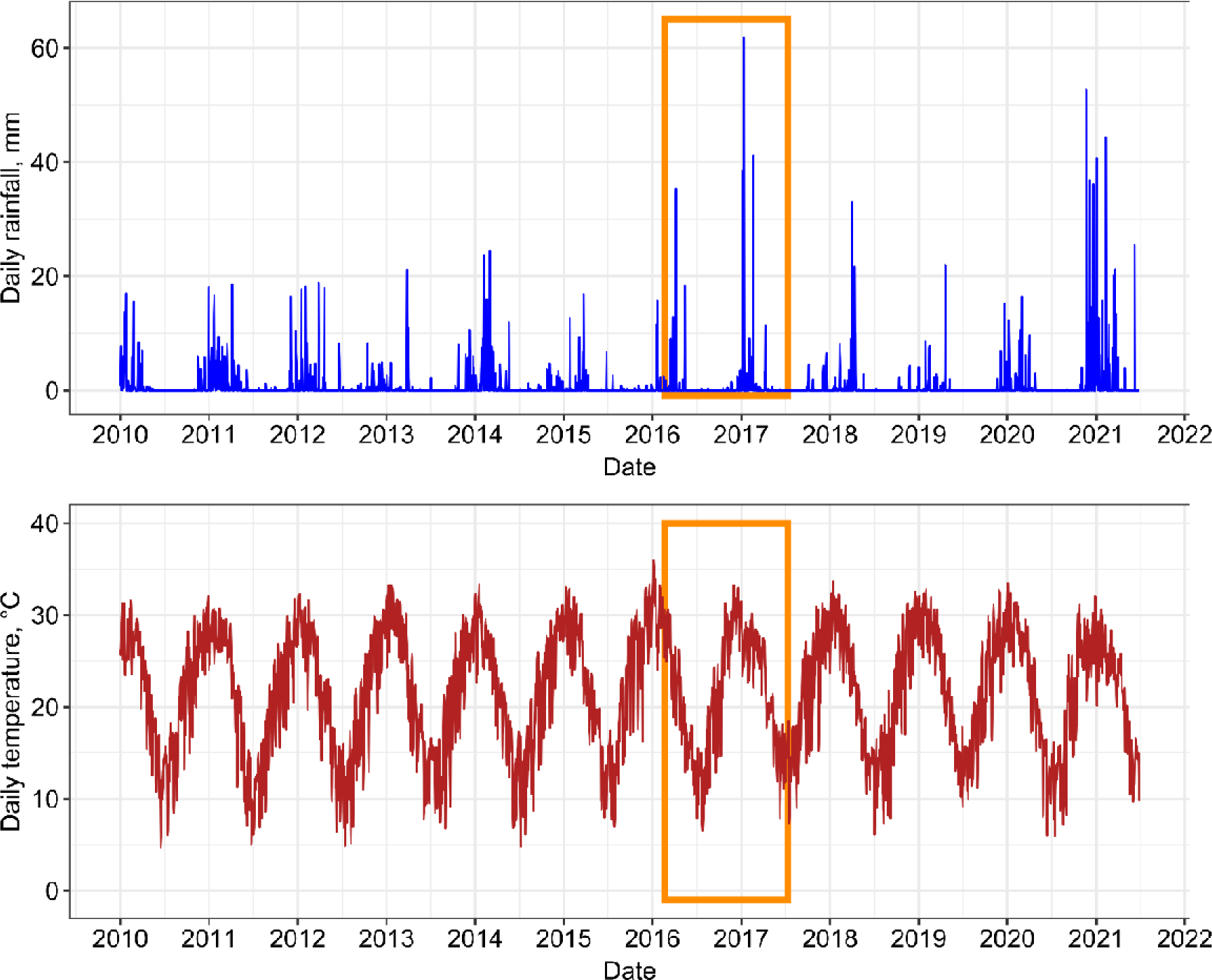
The timing of the experimental pairings in relation to climatic variation at the Kuruman River Reserve. Upper and lower panels document the long-term daily rainfall (mm) and daily average temperature at the field site. The orange rectangle demarcates the period within which experimentally created pairs and established groups (which served as controls) were captured and recaptured. Climate data taken from NASA’s GMAO MERRA-2 assimilation model and GEOS 5.12.4 FP-IT.

**Fig. S11.**
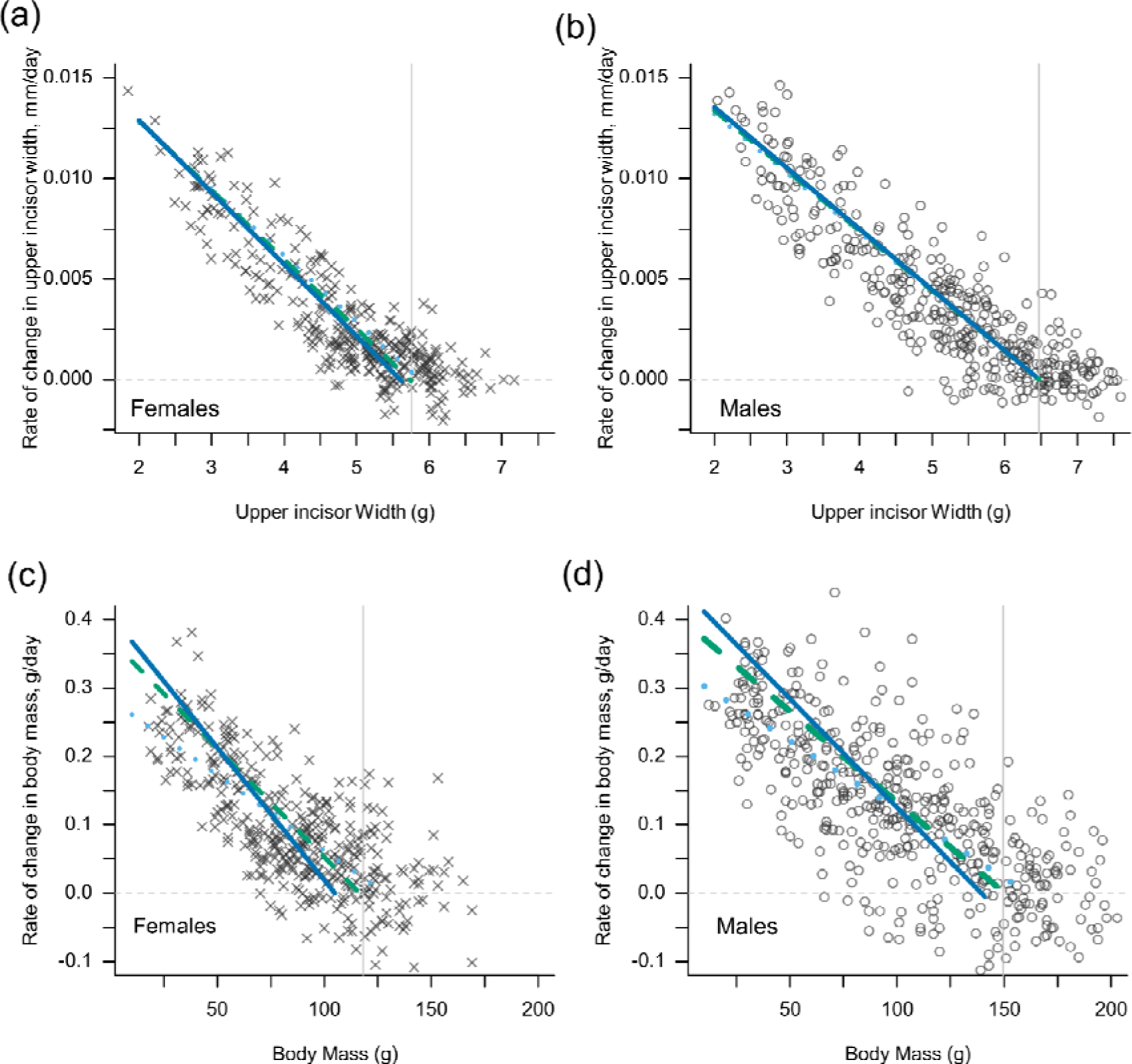
The rate of change in incisor width and body mass for females (a, c) and males (b, d) with variation in initial size and group size. Each point represents the change in incisor width or body mass across two captures - a capture and a recapture. Note that with increasing initial size, the slope of the change in size metric converges on the estimated population-level asymptotes (vertical lines).

### Group size and adult body mass

Because the parameters of the von-Bertalanffy growth curve covary with one another (as parameterised), it is not possible to dissociate the effect of group size on the growth rate constant, *k*, from the effect of group size on asymptotic mass, *A*; *k* itself contributes to the estimation of *A*. We therefore carried out additional analyses to clarify the effect of group size on adult body mass.

For all the males and females whose growth was modelled by the interval equations (Fig. S7), we extracted weight information for all the times that they were captured at least one year after first being captured at less than 1 year of age (< 100g for males, < 80g for females). The weights information therefore represented the body mass of all non-breeding individuals captured beyond 1 year of age, with most data coming from individuals far older than this. This generated a data set of 152 weights from 73 males, and 111 weights from 65 females. At the same time, we calculated the average group size that each of these individuals had experienced in their first year life; for some individuals we had only one value across this period, for others we had up to 3 measures.

The mass of individuals in adulthood of either sex was then modelled according to their current group size and the group size that they experienced in early life (< 1 year of age). The two terms were not strongly correlated (r = 0.26 in males, r = 0.04 in females), and in both the male and the female data sets the variance inflation factor of the two terms was < 1.07. Both early life group size and adult group size could therefore be fitted together in the same model. Adult body mass was fitted in a linear mixed effects model (Gaussian error), with average group size in the first year specified as a continuous variable, and group size in adulthood specified as a categorical variable with three levels: small, medium, and large, separated into the tertiles in each data set. For males at 1-9 (small), 10-16 (medium), >16 (large); for females at 1-7 (small), 8-15 (medium), >15 (large). An additional variable noting the quarterly period of the year was also included (Jan-Mar; Apr-Jun; Jul-Sep; Oct-De). Group size in early life was standardised prior to model fitting, and a random effect of individual identity, nested within group identity, was specified in each case. Models were fitted in the *lme4* package in R. Model outputs are presented in tables S6, and visualised in Fig. S11.

**Fig. S12.**
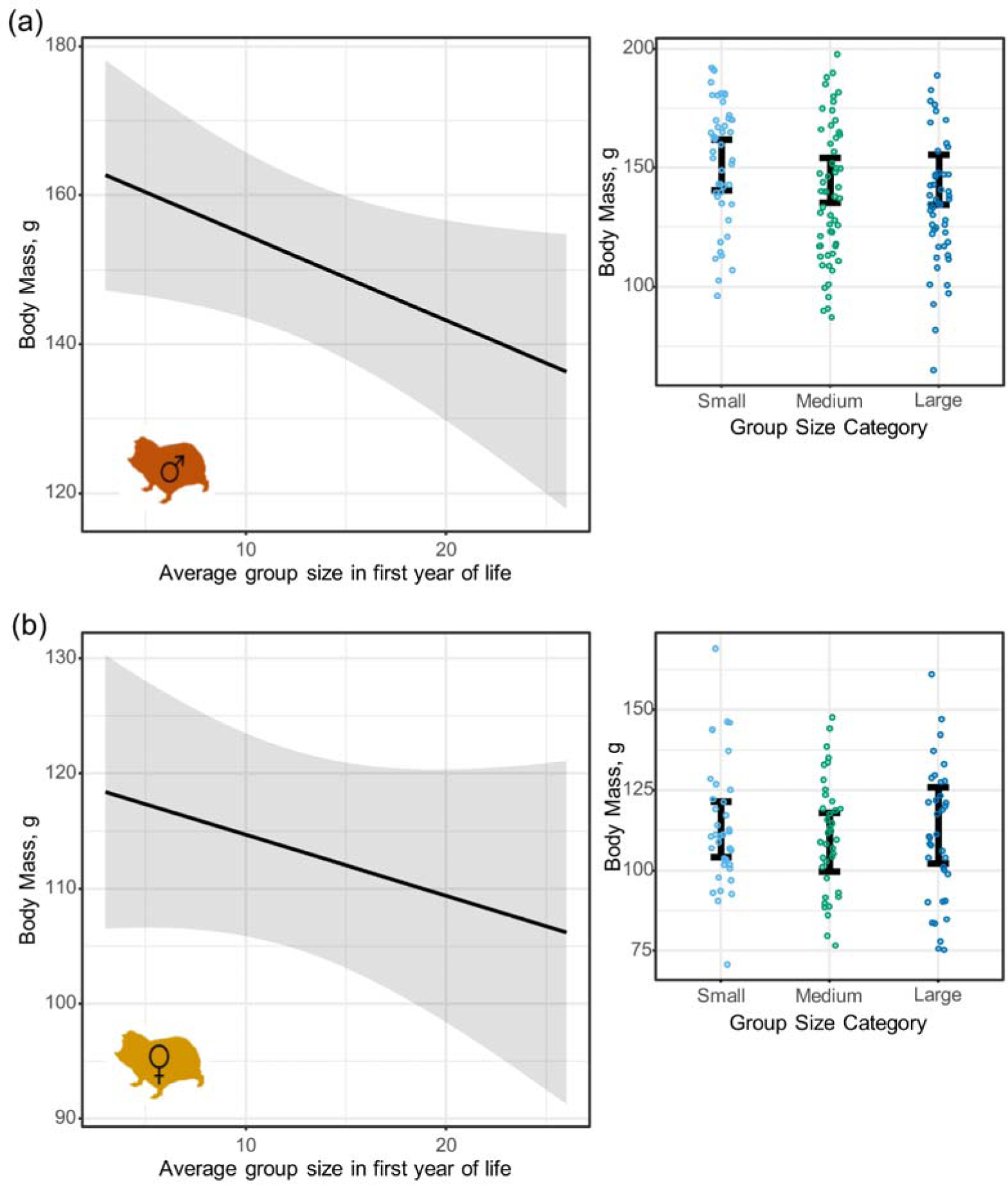
Adult body mass of non-breeding male (a) and female (b) Damaraland mole-rats. Left panels display the predicted effect of average group size in the first year of life on adult mass (< 100g for males, < 80g for females), as estimated from linear mixed effects models (table S11). In both cases, increases in group size during early life reduces body mass in adulthood. By contrast, current group size in adulthood has no effect on adult body mass, as displayed in the right panels. Points denote the raw data across all individuals, with error bars denoting the predicted marginal effects across three group size categories. Group size categories were separated at the tertiles within each data set; for males at 1-9 (small), 10-16 (medium), >16 (large); for females at 1-7 (small), 8-15 (medium), >15 (large). Post-hoc tests performed on the linear mixed effects models found that in neither sex did group size in adulthood have a significant effect on adult body mass (p > 0.46 in all contrasts). Note that group size in adulthood was categorised largely for the purpose of visualisation; treating the term as a continuous variable did not qualitatively affect the results.

**Table S2.**
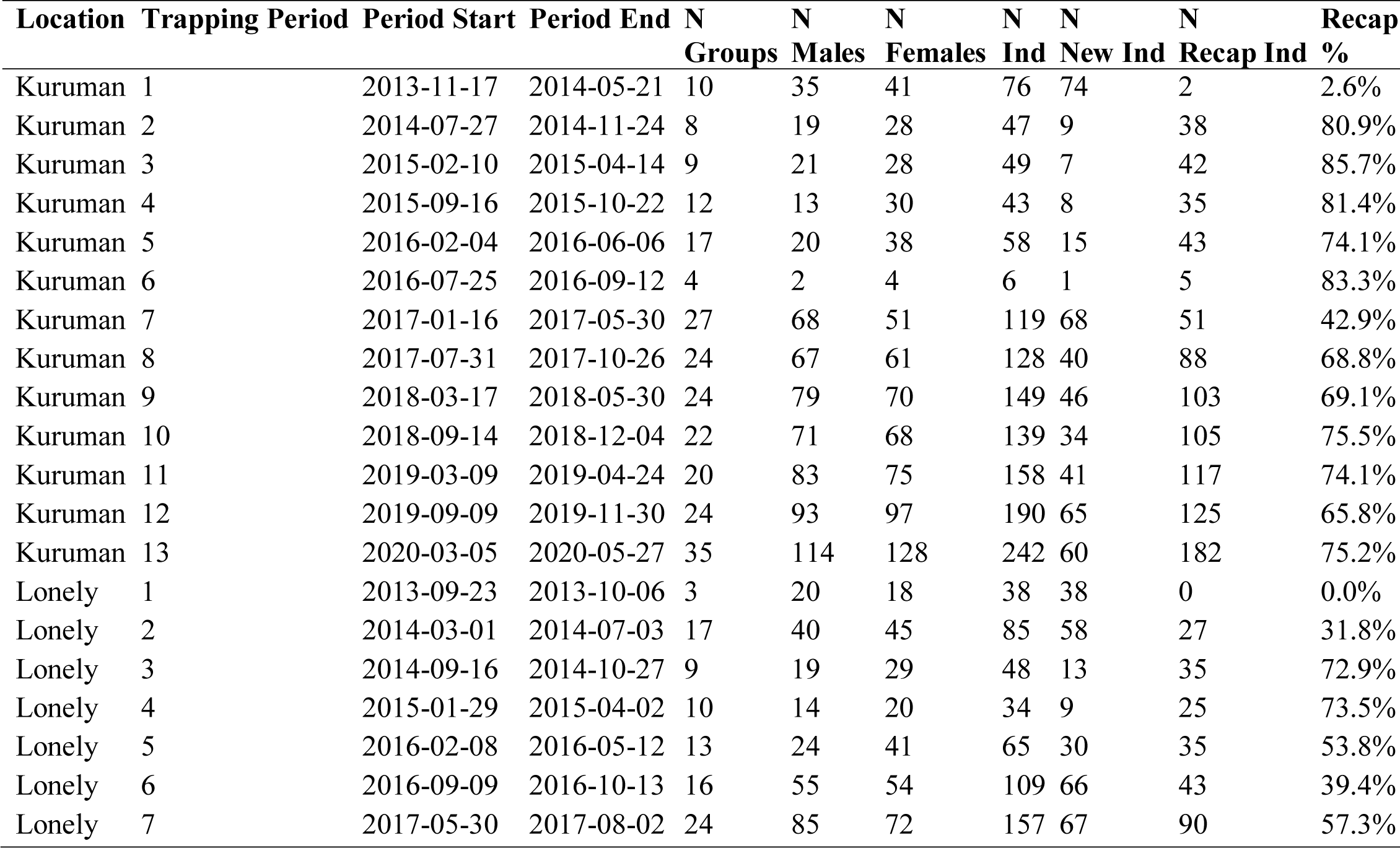
Sum of captures at the Kuruman River Reserve for each trapping period. Recapture rate was calculated as the number of recaptured individuals divided by the total number of individuals captured in the same trapping period. By the end of the first trapping period, two individuals had already been recaptured within new groups (hence 2.6 % recapture rate). Mean recapture rate (N recaptures / N captures within a trapping period, excluding the first trapping period) at Kuruman was 73.1 %, and 54.8% at Lonely.

**Table S3.** Summary of the analyses carried out in the paper, with information on the data used.

| Analysis | Data | Total rows of data | Total unique inds | Total unique groups | Included incomplete captures | Method | Comments |
| --- | --- | --- | --- | --- | --- | --- | --- |
| Status-related survivorship in females | All capture events from females, irrespective of age (weight) at first capture. Additional rows generated for when individuals disappeared or were right-censored.<br>“Fates_Allfemales.csv” | F: 1315 (953 captures) | F: 362 | 80 | Yes | Multi-state Markov model. | - Right-censored individuals that were still present for the last trapping window at either study location.<br>- Four states: i- *in-group non-breeding female*, ii- *single female*, iii- *breeding female*, iv - *disappeared/dead*<br>- Fig. 2. Table S4. |
| Natal dispersal | “FieldMR_Femalephilopatry.csv”<br>“FieldMR_Malephilopatry.csv” | 1571 (1375 captures) | F: 227<br>M: 272 | 56 | Yes | Two-state Markov model | - Right-censored individuals that were still present for the last trapping window at either study location.<br>- Two states: i- *in-group non-breeder*, ii- *disappeared/dead/known disperser*<br>- Fig.s S7, S8. Table S1. |
| Growth: body mass | Body mass across successive capture events (a capture and recapture)<br>“FieldMR_Growth_BodyMass.csv” | F: 381<br>M: 456 | F: 193<br>M: 214 | F: 48<br>Male: 39 | NA | Non-linear mixed effects model with von Bertalanffy interval equation. Males and females analysed separately. | - Removed breeding females so that pregnancy weights were excluded.<br>- Only chose recaptures that occurred 100-365 days after the first capture.<br>- Fig. 5, Fig. S11. Table S6<br>- Follow-up analysis to check the effect of group size on asymptotic mass (above; Fig. S12). |
| Growth: incisor width | Incisor width across successive capture events (a capture and recapture)<br>“FieldMR_Growth_TeethWidth.csv” | F: 328<br>M: 381 | F: 180<br>M: 198 | F: 47<br>Male: 38 | NA | Non-linear mixed effects model with von Bertalanffy interval equation. Males and females analysed separately. | - Only chose recaptures that occurred 100-365 days after the first capture.<br>- Fig. S11. Table S8. |
| Body condition of | All body mass measurements with an associated skeletal trait measurement | F incisor width: | F incisor width: | F incisor width: | NA | Scaled major axis regression of ln(body | -Included information from females that were either an in-group non-breeder or a |
| females | (either teeth width of body length)<br>“*FieldMR_SingleFemaleCondition_BodyLength.csv*”<br>“*FieldMR_SingleFemaleCondition_TeethWidth.csv*” | 353<br>F body length: 419 | 177<br>F body length: 213 | 64<br>F body length: 67 |  | mass) ~ ln(skeletal trait).<br>In both cases, a separate slope of intercept for single females was not supported. | single female. As the main question was whether single females had a similar body condition to in-group non-breeding females, we only considered in-group non-breeders when their skeletal trait was greater than the smallest recorded value in single females (i.e. removing young females still in their natal group to ensure this didn't bias allometric scaling).<br>- Fig. 3. |
| Within-group recruitment rate (longitudinal) | All complete group captures and recaptures separated by 100-365 days.<br>“*FieldMR_Recruitment_Longitudinal.csv*” | 78 capture-recapture events | NA | 33 groups | No | Generalised linear mixed effects model with Poisson error distribution. Included offset term for time interval. | -Within-group recruits classed as all new individuals in the group on recapture that were less than year of age (body mass < 80g for females, body mass < 100g for males”)”<br>- Only included groups if capture history indicated the presence of a breeding female.<br>- Fig. 4. Table S5. |
| Within-group recruitment rate (experimental) | The number of new recruits on the first recapture of experimentally created pairs, which were compared to the number of recruits in established groups captured across the same time period.<br>“*FieldMR_Recruitment_Experimental.csv*” | Pairings: 8<br>Time-matched groups: 9 | NA | NA | No | t-test on recruitment rate | - Fig. 4. |

**Table S4.**
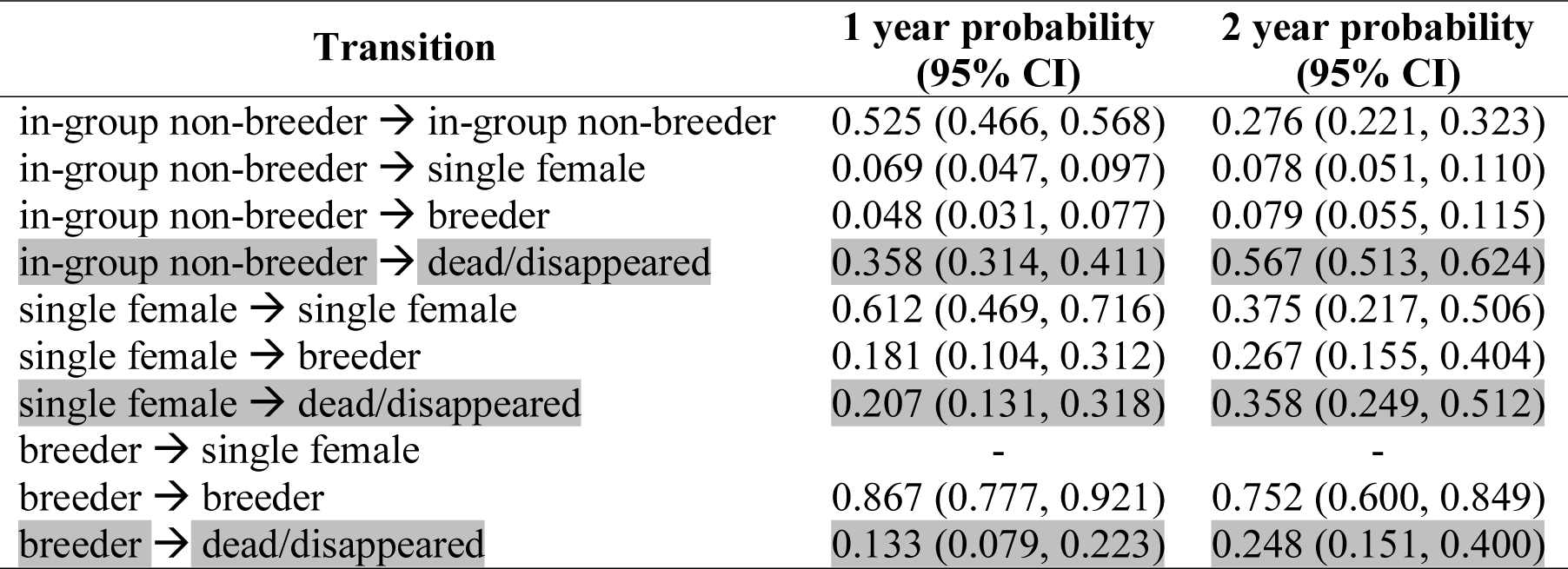
Status-related survivorship. Probability of transitioning between states over a one-year and a two-year period. Transition probabilities estimated from a multi-state model fitted to all females that were captured across the duration of the study. n = 362 females. Estimates related to survival probability are filled in grey. Confidence intervals estimated using the delta method.

| Transition | 1 year probability<br>(95% CI) | 2 year probability<br>(95% CI) |
| --- | --- | --- |
| in-group non-breeder → in-group non-breeder | 0.525 (0.466, 0.568) | 0.276 (0.221, 0.323) |
| in-group non-breeder → single female | 0.069 (0.047, 0.097) | 0.078 (0.051, 0.110) |
| in-group non-breeder → breeder | 0.048 (0.031, 0.077) | 0.079 (0.055, 0.115) |
| in-group non-breeder → dead/disappeared | 0.358 (0.314, 0.411) | 0.567 (0.513, 0.624) |
| single female → single female | 0.612 (0.469, 0.716) | 0.375 (0.217, 0.506) |
| single female → breeder | 0.181 (0.104, 0.312) | 0.267 (0.155, 0.404) |
| single female → dead/disappeared | 0.207 (0.131, 0.318) | 0.358 (0.249, 0.512) |
| breeder → single female | - | - |
| breeder → breeder | 0.867 (0.777, 0.921) | 0.752 (0.600, 0.849) |
| breeder → dead/disappeared | 0.133 (0.079, 0.223) | 0.248 (0.151, 0.400) |

**Table S5.**
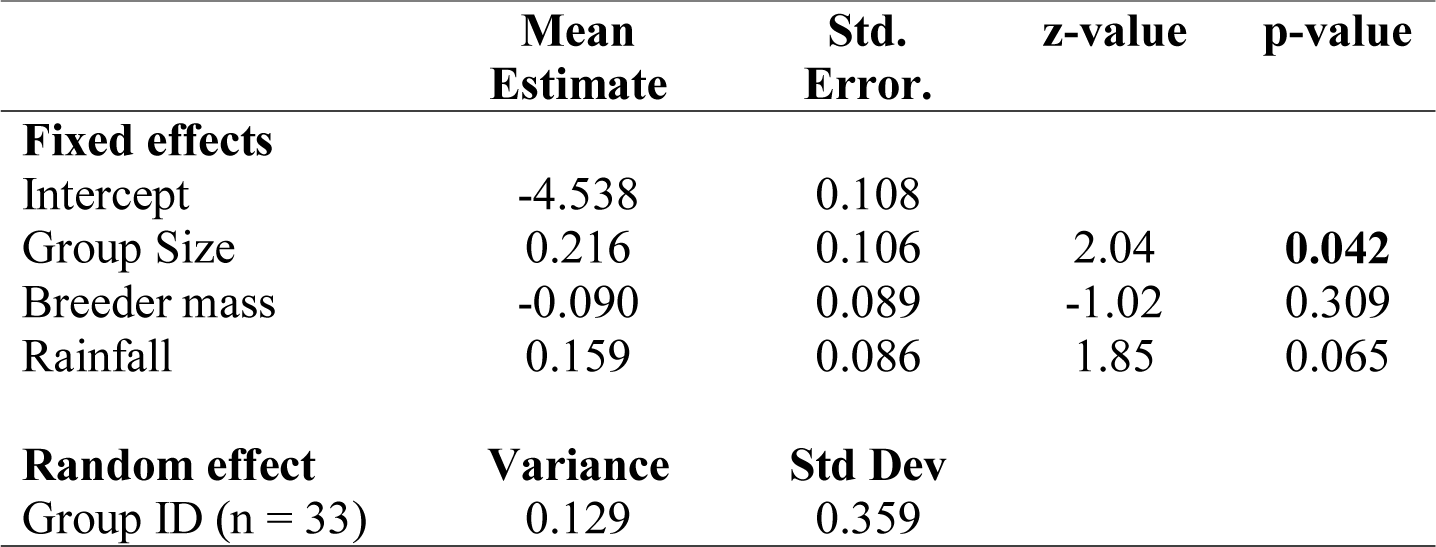
The recruitment rate of offspring into groups in Damaraland mole-rats in the wild. Recruitment was modelled with a generalised linear model with negative binomial error distribution. Models were fitted to n = 77 group-level recapture events in 33 distinct breeding groups. All estimates are provided on the link scale (log-link), and continuous variables are z-score transformed. Models also included an offset for the log(time between capture and recapture).

**Table S6.** Body mass growth of Damaraland mole-rats in the wild. Model summaries provide estimates from the non-linear mixed effects that incorporated group size into von Bertalanffy interval equations. Models were fitted to n = 381 recapture events in females, and n = 456 recapture events in males. Estimates provide the population mean for the fixed effects, the standard deviation for the random effects.

| *Females* | **Mean**<br>**Estimate** | **Std.**<br>**Error.** | **t-value, df** | **p-value** |
| --- | --- | --- | --- | --- |
| **Fixed effects** |  |  |  |  |
| *A* | 118.26 | 2.01 | 58.72, 185 | < **0.001** |
| *A<sub>GS</sub>* | -9.42 | 1.47 | 6.41, 185 | < **0.001** |
| *K* | 0.00447 | 0.00003 | 16.02, 185 | < **0.001** |
| *k<sub>GS</sub>* | 0.00155 | 0.00032 | 4.90, 185 | < **0.001** |
| **Random effect** | **Variance** | **Residual** |  |  |
| *A* ~ 1\|AnimalID | 15.79 | 9.18 |  |  |
| *k* ~ 1\|AnimalID | Negligible |  |  |  |
| *Males* | **Mean**<br>**Estimate** | **Std.**<br>**Error.** | **t-value, df** | **p-value** |
| **Fixed effects** |  |  |  |  |
| *A* | 149.70 | 2.84 | 52.74, 239 | < **0.001** |
| *A<sub>GS</sub>* | -8.10 | 2.02 | 4.00, 239 | < **0.001** |
| *k* | 0.00368 | 0.00002 | 17.90, 239 | < **0.001** |
| *k<sub>GS</sub>* | 0.00084 | 0.00020 | 4.36, 239 | **0.001** |
| **Random effect** | **Variance** | **Residual** |  |  |
| *A* ~ 1\|AnimalID | 21.47 | 12.51 |  |  |
| *k* ~ 1\|AnimalID | Negligible |  |  |  |

**Table S7.** The adult body mass of mole-rat non-breeders. Female and male mass was modelled using a linear mixed effects model with Gaussian error distribution. In either case, models were fitted to non-breeding individuals that were captured at least one year after they were first captured at less than 1 year of age (< 100g for males, < 80g for females). They therefore represent the average body mass of all non-breeding individuals captured beyond 1 year of age, with most data coming from individuals far older than this. Group size categories were separated at the tertiles within each data set; for males at 1-9 (small), 10-16 (medium), >16 (large); for females at 1-7 (small), 8-15 (medium), >15 (large). n = 65 unique females from 25 groups. n = 73 unique males from 23 groups. See Fig. **S11**.

| *Females (n = 111)* | **Mean**<br>**Estimate** | **Std.**<br>**Error.** | **t-value,**<br>**df** | **Chisq, df** | **p-value** |
| --- | --- | --- | --- | --- | --- |
| **Fixed effects** |  |  |  |  |  |
| Intercept | 112.95 | 4.41 |  |  |  |
| Adult group size: medium | -4.12 | 4.62 | -0.892 | 1.23, 2 | 0.54 |
| Adult group size: large | 1.15 | 6.27 | 0.183 |  |  |
| Yearly quarter: second | -4.55 | 3.87 | -1.178 | 1.90, 3 | 0.59 |
| Yearly quarter: third | -4.15 | 4.11 | -1.009 |  |  |
| Yearly quarter: fourth | -2.06 | 3.97 | -0.511 |  |  |
| Early life group size | 3.08 | 2.62 | -1.176 | 1.23, 1 | 0.27 |
| **Random effect** |  |  |  |  |  |
|  | **Variance** | **Std Dev** |  |  |  |
| Animal ID:Group ID | 162.2 | 12.73 |  |  |  |
| Animal ID | 114.6 | 10.71 |  |  |  |
| Residual | 128.5 | 11.34 |  |  |  |
| **Males (n = 152)** |  |  |  |  |  |
|  | **Mean**<br>**Estimate** | **Std.**<br>**Error.** | **t-value,**<br>**df** |  |  |
| **Fixed effects** |  |  |  |  |  |
| Intercept | 151.30 | 5.44 |  |  |  |
| Adult group size: medium | -6.39 | 4.49 | -1.423 | 2.13, 2 | 0.34 |
| Adult group size: large | -6.15 | 5.36 | -1.148 |  |  |
| Yearly quarter: second | -14.22 | 3.36 | -4.238 | 27.15, 3 | <0.001 |
| Yearly quarter: third | -15.53 | 3.24 | 4.789 |  |  |
| Yearly quarter: fourth | -12.03 | 3.57 | -3.373 |  |  |
| Early life group size | -6.56 | 3.33 | -1.967 | 2.99, 1 | 0.084 |
| **Random effect** |  |  |  |  |  |
|  | **Variance** | **Std Dev** |  |  |  |
| Animal ID:Group ID | 493.2 | 22.21 |  |  |  |
| Animal ID | 159.4 | 12.62 |  |  |  |
| Residual | 141.0 | 11.88 |  |  |  |

**Table S8.**
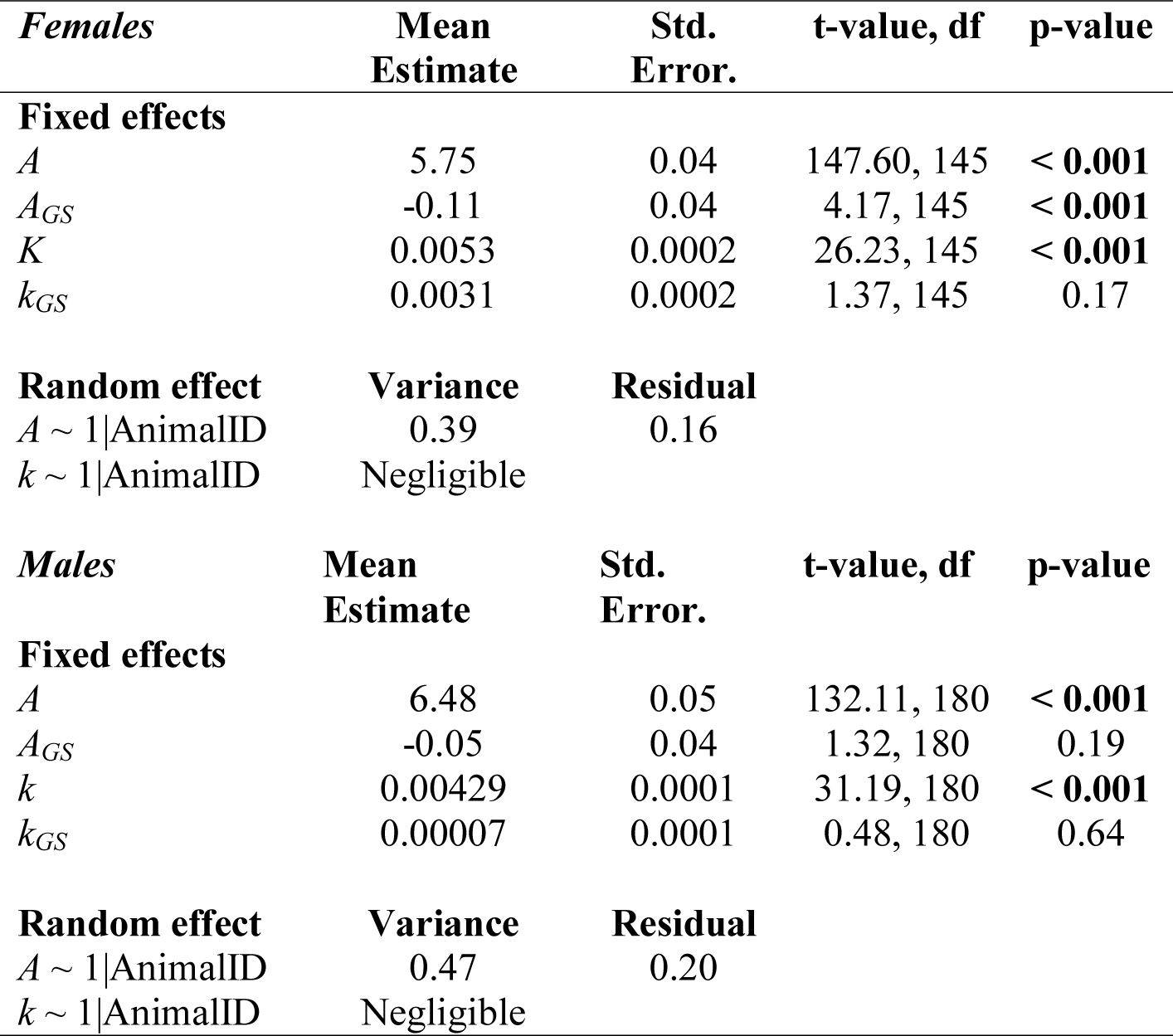
Incisor width growth of Damaraland mole-rats in the wild. Model summaries provide estimates from the non-linear mixed effects that incorporated group size into von Bertalanffy interval equations. Models were fitted to n = 328 recapture events in females, and n = 381 recapture events in males. Estimates provide the population mean for the fixed effects, the standard deviation for the random effects.

